# HDAC4 Inhibits NMDA Receptor-Mediated Stimulation of Neurogranin Expression

**DOI:** 10.1101/2024.07.09.602724

**Authors:** Raquel de Andrés, Elena Martínez-Blanco, F. Javier Díez-Guerra

## Abstract

The coordination of neuronal wiring and activity within the central nervous system (CNS) is crucial for cognitive function, particularly in the context of aging and neurological disorders. Neurogranin (Ng), an abundant forebrain protein, modulates calmodulin (CaM) activity and deeply influences synaptic plasticity and neuronal processing. This study investigates the regulatory mechanisms of Ng expression, a critical but underexplored area for combating cognitive impairment. Utilizing both in vitro and in vivo hippocampal models, we show that Ng expression arises during late developmental stages, coinciding with synaptic maturation and neuronal circuit consolidation of. We observed that Ng expression increases in neuronal networks with heightened synaptic activity and identified GluN2B-containing N-methyl-D-aspartate (NMDA) receptors as key drivers of this expression. Additionally, we discovered that nuclear-localized HDAC4 inhibits Ng expression, establishing a regulatory axis that is counteracted by NMDA receptor stimulation. Analysis of the Ng gene promoter activity revealed regulatory elements between the −2.4 and −0.85 Kbp region, including a binding site for RE1-Silencing Transcription factor (REST), which may mediate HDAC4’s repressive effect on Ng expression. Further analysis of the promoter sequence revealed conserved binding sites for the myocyte enhancer factor-2 (MEF2) transcription factor, a target of HDAC4-mediated transcription regulation. Our findings elucidate the interplay between synaptic activity, NMDAR function, and transcriptional regulation in controlling Ng expression, offering insights into synaptic plasticity mechanisms and potential therapeutic strategies to prevent cognitive dysfunction.

## Introduction

Human cognitive abilities are intimately linked to the formation and modulation of interconnected neuronal networks through mechanisms collectively known as synaptic plasticity. These mechanisms are responsible for the strengthening or weakening of synaptic connections in response to activity patterns. Calcium (Ca2+)-mediated signaling plays a pivotal role in the mechanisms underlying synaptic plasticity^1^. Calmodulin (CaM), a ubiquitous calcium-binding protein, regulates multiple proteins, including kinases, phosphatases, ion channels and neurotransmitter receptors^2^, thereby regulating various forms of synaptic plasticity, such as short-term plasticity, Hebbian plasticity (long-term potentiation (LTP) and long-term depression (LTD)), homeostatic plasticity, and synaptic remodeling^3^. Rapid adaptive plasticity typically involves modifications of existing proteins or lipids, while more enduring plasticity requires gene transcription and new protein synthesis. Activity-regulated gene expression affects genes encoding synaptic proteins that modulate synaptic transmission^4^, thereby influencing neuronal network functions.

Neurogranin (Ng) is a small protein primarily expressed in telencephalic areas, including the amygdala, cerebral cortex, hippocampus and striatum^5^. It is highly expressed in principal neurons, where it is localized in soma, dendrites and dendritic spines^6^. In the rat brain, Ng levels peak in the third postnatal week, coinciding with a period of active synaptogenesis, and remain elevated throughout adulthood^7^. Ng binds to CaM and to phosphatidic acid (PA)^8^ through a central IQ motif that can be phosphorylated by PKC. Ng has a higher affinity for apo-CaM than for Ca^2+^/CaM. Its phosphorylation or high Ca^2+^ levels prevent Ng from binding to CaM. Given Ng’s high intracellular levels^9^, it is believed that most apo-CaM is sequestered by Ng in neurons. This would limit CaM activation at low Ca^2+^ levels^10^ and regulate CaM availability to its targets, thus influencing synaptic plasticity^11^.

Ng content of human brain decreases with age and in diseases associated to cognitive impairment^12^. Cerebrospinal fluid (CSF) from patients with mild cognitive impairment (MCI) shows elevated levels of a Ng peptide^13,14^, supporting Ng as an early biomarker of cognitive decline^15^. Ng knockout (KO) mice show impaired hippocampal-dependent learning and memory tasks^16^, whereas increased Ng expression improves cognitive function^17,18^. In cultured neurons, Ng expression increases spine number and neuronal maturity^19^, while in vivo enhances PSD-95^18^ and phosphorylated CaMKII levels in the CA1 region of the hippocampus^20^. Additionally, increased Ng expression rescues amyloid-β (Aβ)-induced synaptic deficits and restores synaptic transmission through its ability to bind CaM and to activate CaMKII^21^.

In this study, we investigated the regulation of Ng expression in both in vitro and in vivo models. Using organotypic hippocampal slice cultures and adult neurogenesis in the Dentate Gyrus (DG), we demonstrate that Ng expression develops at late stages of neuronal maturation, periods marked by intense synaptogenesis and integration into neuronal networks. Our in vitro experiments show that N-methyl-D-aspartate (NMDA) receptor (NMDAR) signaling is essential for driving Ng expression. Culture media formulations that enhance endogenous synaptic activity greatly increased Ng levels, an effect blocked by NMDAR antagonists targeting the GluN2B subunit. We found that the histone deacetylase HDAC4 plays a repressive role that can be antagonized by NMDAR activation. Finally, we analyzed the transcriptional activity of various regions of the Ng gene promoter using a fluorescent reporter. We propose that synaptic activation of GluN2B-containg NMDARs triggers signaling pathways that fine-tune Ng expression. These findings advance our understanding of Ng expression regulation and pave the way for strategies aimed at upregulating Ng to improve cognitive performance in aging and early stages of neurodegenerative diseases.

## Materials and Methods

### Animals and ethics compliance

Wistar rats and C57BL/6 mice were bred at the animal facility of the Centro de Biología Molecular Severo Ochoa (CBMSO). All procedures were carried out in accordance with the Spanish Royal Decree 1201/2005 for the protection of animals used in scientific research and the European Union Directive 2010/63/EU regarding the protection of animals used for scientific purposes. The procedures were approved by the institutional and regional local ethics committees.

### Reagents

Fetal bovine serum (FBS), Dulbecco’s modified Eagle’s medium (DMEM), 0.25% trypsin, Neurobasal, Neurobasal plus, B27, and B27+ media and supplements were obtained from Thermo Fisher Scientific. The protease inhibitor cocktail was from Biotools (B14001). Total protein was measured using the Bradford Protein Assay kit (Bio-Rad). Pre-stained protein markers VI (10-245 kDa) was from PanReac-AppliChem. Immobilon-P membranes and ECL western blotting reagent was from Millipore. Oligonucleotides were purchased from Integrated DNA Technologies (IDT). 1-beta-arabino-furanosylcytosine (AraC #251010) was from Calbiochem (#251010). D-2-amino-5-phosphonovalerate (AP5), memantine hydrochloride, N-methyl-D-aspartic acid (NMDA), (+)-MK-801 maleate, tetrodotoxin (TTX), 2,3-Dioxo-6-nitro-1,2,3,4-tetrahydrobenzo[f]quinoxaline-7-sulfonamide disodium salt (NBQX), ifenprodil, (-)-bicuculline methiodide (BIC), and HDAC inhibitors M275 (#6208), Trichostatin A (#1406), LMK 235 (#4830) and MC 1568 (#4077) were from Tocris. UBP714 (NMDAR PAM, # HB8161) was from Hello Bio. Compound 8 (C8) was obtained from Dr. H. Bading lab. Paraformaldehyde (PFA) was from Merck. Antibodies and plasmids used are listed in Supplementary Table 1.

### Primary cultures of rat hippocampal neurons

Cultures of rat hippocampal neurons were prepared as previously described^22^. Embryos at 19-20 days gestation were obtained and kept in chilled incomplete Hank’s medium during dissection. The brains were extracted, and hippocampi dissected, with meninges carefully removed. Hippocampi were washed five times in incomplete Hank’s, followed by a 0.25% trypsin treatment at 37°C for 15 minutes to aid cell dissociation. After removing trypsin with two washes in incomplete Hank’s, tissue was transferred to complete Hank’s with DNase (0.04 mg/ml) for mechanical dissociation using Pasteur pipettes and 22G needles. The dissociated cells were filtered through a 70 µm nylon mesh, centrifuged at 1200 rpm for 5 minutes, resuspended in plating medium, and counted. Neurons were plated on culture dishes pre-treated with 0.1 mg/ml Poly-L-lysine (PLL) in borate buffer at a density of 25,000 cells/cm², and on coverslips pre-treated with 0.25 mg/ml PLL in borate buffer (pH 8.0) at a density of 12,000 cells/cm². Coverslips were sterilized with 65% nitric acid and heat (180°C) before PLL treatment. After 3 hours of incubation to allow cell adhesion, the plating medium was replaced with Neurobasal supplemented with B27 and GlutaMAX. On day 3 in vitro (DIV3), 1µM AraC (1-β-arabinofuranosylcytosine) was added to eliminate glial cells and obtain pure neuron cultures. At DIV7, half of the medium was replaced with fresh Neurobasal + B27 (NB) or Neurobasal Plus + B27+ (NB+). Cultures were maintained at 37°C with 5% CO2 and processed during the third week in vitro.

### Organotypic cultures of rat hippocampal slices

All procedures were under sterile conditions. Rat pups aged 5-7 days (P5-7) were anesthetized and decapitated. The brains were immersed in cold dissection medium (KCl 4 mM, NaHCO3 26 mM, MgCl2 5 mM, CaCl2 1mM, Glucose 10mM and sacarose 233.7mM), their hippocampi extracted and cut into 400 µm thick coronal sections using a chopper. The slices were placed on inserts with porous nitrocellulose membranes and maintained in organotypic culture medium (MEM, FBS 20%, Glutamine 1 mM, CaCl2 1 mM, MgSO4 5 mM, insulin 1mg/L, ascorbic acid 0,0012%, NaHCO3 5.2 mM, HEPES 30 mM and glucose 13mM) at 35.5°C and 5% CO2. The medium was replaced every 3 days.

### Preparation of viral vectors in HEK-293T cells

HEK-293T cells were cultured in DMEM supplemented with 10% FBS (Gibco) and maintained at 37 °C with 5% CO2. Passaging was performed twice a week.

#### Lentivirus production

All lentiviral constructs derived from the vector pLOX-Syn-DsRed-Syn-GFP^23^, kindly donated by Dr. FG Scholl. HEK-293T cells were seeded 24 hours before transfection at a density of 3 million cells per p100 plate in DMEM with 10% FBS to achieve 80% confluency at transfection time. 24 hours later, the medium was replaced with OptiMEM (Gibco). The transfection mix (8µg of the packaging plasmid, 4µg of pCMV δR8.74, 2µg of pMD2.G, and 28µg PEI Max (Polysciences) in OptiMEM) was incubated for 20 minutes at room temperature (RT) and then added to the cells. 5 hours later, the transfection medium was replaced with DMEM. After 48 hours, the medium containing lentiviral particles was collected, filtered through a 0.45 µm filter, aliquoted, and stored at −80°C until use. Neuronal cultures were infected with lentiviral particles at DIV3.

#### Adeno-associated virus (AAV) production

All AAV constructs derived from the original vector pAAV_hsyn_NES-his-CaMPARI2-F391W-WPRE-SV40, Addgene #101061. HEK-293T cells were seeded 24 hours before transfection at a density of 7 million cells per p150 plate in DMEM with 10% FBS to achieve 70% confluency at transfection. The next day, the medium was replaced with DMEM + 5% FBS. The transfection mix included 12.5µg of the AAV plasmid, 25µg of pFΔ6, 6.25µg of pRV1, 6.25µg of pH21 (helper, capsid, replication^24^), 130.5µl of 2M CaCl2, and 900µl of 2X HBS in a final volume of 1.8ml. After a 20-minute incubation at room temperature, the mix was added to the cells. Sixteen hours later, the medium was replaced with fresh DMEM + 10% FBS. After 48 hours, the medium was discarded, the cells collected in PBS and lysed using 0.4% sodium deoxycholate and 50 U/ml benzonase. Lysates were purified on heparin columns (GE Healthcare Life Sciences) and concentrated using 100,000 Da concentrator filters (Millipore). The concentrate was filtered (0,22 µm), aliquoted, and stored at −80°C until use. Neuronal cultures were infected with AAV particles at DIV3 using only half the normal volume of medium. The other half was stored at 4°C. The next day, the medium with AAV particles was removed and replaced with a 1:1 mixture of the stored medium and fresh Neurobasal medium.

### Design and production of shRNAs

Using SiDirect 2.0 (http://sidirect2.rnai.jp), targets for HDAC4 silencing were selected based on the highest shRNA stability and the fewest off-targets. Two target sequences were chosen in untranslated regions (UTRs) of rat HDAC4 mRNA: 121-143 (5’ UTR) and 3923-3945 (3’ UTR). Complementary oligonucleotides with sense and antisense sequences were designed, hybridized and cloned into the pLKO Tet-ON vector, where shRNAs are controlled by the H1 promoter, and the expression dependent on doxycycline treatment. As a control we used pLKO.1-Scrambled (Addgene #136035). The sequences of the shRNAs are the following:

Fw-HDAC4-shRNA_121-143_:

5’-CCGGTGATTACTTGGTTTACAAGACGGCTTCCTGTCACTCTTGTAAACCAAGTAATCCATTTTTTG-3’

Rv-HDAC4-shRNA_121-143_

5’-AATTCAAAAAATGGATTACTTGGTTTACAAGAGTGACAGGAAGCCGTCTTGTAAACCAAGTAATCA-3’

Fw-HDAC4-shRNA_3923-3945_:

5’-CCGGTGCTTAACATTTGATTCTTACCGCTTCCTGTCACTAAGAATCAAATGTTAAGCACTTTTTTG-3’

Rv-HDAC4-shRNA_3923-3945_:

5’-AATTCAAAAAAGTGCTTAACATTTGATTCTTAGTGACAGGAAGCGGTAAGAATCAAATGTTAAGCA-3’

### Preparation of brain sections for immunostaining

C57BL/6 mice were anesthetized with xylazine (10 mg/kg) and ketamine (80 mg/kg) and perfused via the left ventricle, with 0.9% NaCl to clear the blood, followed by 4% paraformaldehyde (PFA) in phosphate buffer (PB) pH 7.4. The right atrium was cut to allow blood exit. Then, the brain was removed and immersed in the fixative solution for 24 hours at 4°C with gentle agitation. After extensive washing in PB, the brains were embedded in 4% agarose/10% sucrose and sectioned at 50 µm using a vibratome (Leica VT1200S). Sections were stored at −20°C in a glycol mixture until use. Before staining, sections were washed with PB to remove glycol, then incubated for 48-72 hours at 4°C with primary antibodies in a blocking/permeabilization solution containing 1% bovine serum albumin (BSA) and 0.1% Triton-X 100 (TX-100) in PB. After washing with blocking/ permeabilization solution, sections were incubated with secondary antibodies (Alexa Fluor conjugated, Thermofisher) in PB for 2 hours at room temperature. In some cases, amplification was performed using biotinylated secondary antibodies and streptavidin-Alexa594. Sections were then incubated with DAPI (0,2 µg/mL) for 5 minutes to stain nuclei. Finally, sections were washed in PB for several hours before mounting on gelatin-coated slides with Mowiol.

### Immunofluorescence of Cell Cultures

Coverslips with cells were briefly washed with PBS and then fixed with 4% PFA in PBS for 15 minutes at room temperature (RT). The fixative was removed, and cells were incubated with 0.2M glycine (pH 8) for 10 minutes at RT to quench residual PFA. After two PBS washes, cells were incubated for 30 minutes in a blocking/permeabilization solution containing 1% BSA, 0.1% TX-100, and 1% horse serum. Primary antibodies were incubated overnight at 4°C in a solution of 1% BSA and 1% horse serum. The next day, after three 10-minute washes with PBS, cells were incubated with secondary antibodies in PBS for 1 hour at RT. After nuclear staining with DAPI, cells were washed three more times and mounted on slides with Mowiol.

### In Vivo EdU Labeling

For labeling mitotic cells in the adult neurogenesis model, 5-Ethynyl-2′-deoxyuridine (EdU) was injected intraperitoneally into wild-type and fractalkine receptor deficient (CX3CR1 -/-) 2-month-old mice, in 5 doses of 35 mg/kg over 3 consecutive days. Mice were perfused weekly from the 3rd to the 8th week post-injection.

### EdU Detection via Click Reaction

After perfusion and brain sectioning as described above, the tissue was washed in PB and permeabilized in PB containing 0.1% TX-100 for 20 minutes with agitation. Non-specific binding was then blocked for 1 hour with agitation in a solution containing 0.1% TX-100 and 1% BSA in PB. The tissue was incubated with primary antibodies for 72 hours at room temperature (RT), followed by extensive washing in PB to remove residual antibodies. Subsequently, the tissue was incubated with secondary antibodies for 1.5 hours. In some cases, amplification was performed using biotinylated secondary antibodies and streptavidin-Alexa594. Before the click reaction, the tissue was incubated in a solution containing PB, 3% BSA and 0.1% TX-100 to reduce background signal. For the click reaction, tissue sections were immersed and incubated for 30 minutes in the dark in the following mixture: 100 mM Tris HCl, 2 mM CuSO4, 10 µM Alexa Fluor 488 Azide, and 100 mM ascorbic acid. Then, the sections were incubated counterstained DAPI. Following extensive washes in PB, the sections were mounted on gelatin-coated slides with Mowiol.

### Protein Extraction and Western Blot

Cell extracts were prepared using a buffer containing: NaCl 50 mM, EDTA 1 mM, DTT 2 mM, TX-100 0.5%, Tris-HCl 25 mM (pH6.8), and a cocktail of protease and phosphatase inhibitors. Homogenization was done using a 23G needle (20 passes), followed by centrifugation at 17,500 xg for 20 minutes at 4°C, and the supernatants collected. For brain extracts, tissue was weighed and homogenized with 5 volumes of the same extraction buffer using needles of decreasing gauge sizes. The homogenates were centrifuged at 17,500 xg for 20 minutes at 4°C, and the supernatants collected. Protein concentration was determined using the Bradford method (Bio-Rad). 10-15 µg of total protein per sample was loaded onto SDS-PAGE lanes and separated by molecular weight at a constant voltage of 120V (2.5 hours). Proteins were then transferred to PVDF membranes (Millipore) using a semi-dry transfer system at a constant current of 400mA for 30 minutes in transfer buffer (Tris 22.5 mM, glycine 170mM, methanol 20%) Membranes were blocked in 5% non-fat milk in TBS-T (0.05% Tween-20) for 45 minutes at room temperature with agitation and incubated with primary antibodies in TBS-T overnight at 4°C with agitation. After three 10-minute washes in TBS-T, membranes were incubated with HRP-conjugated secondary antibodies in TBS-T for 1 hour at room temperature with agitation. Finally, membranes were washed three times with TBS-T and once with TBS. Protein bands were detected using ECL reagent (Millipore) and the Amersham Imager 680 (GE Healthcare Life Sciences). Densitometric analysis of protein bands was performed using ImageJ software.

### RT-PCR

Total RNA was extracted using TRIzol reagent (Invitrogen) following the manufacturer’s instructions. Subsequently, RNA was quantified at 260nm absorbance, and equal amounts of RNA were reverse-transcribed into DNA using the First-Strand cDNA Synthesis Kit (NZYTech). A conventional PCR was performed with 1ng of each sample to amplify Ng transcripts. GAPDH was used as a reference. The following oligonucleotides were used for PCR amplification:

Fw-Ng: 5’-GACTGCTGCACGGAGAGC-3’

Rv-Ng: 5’-CAGTTCTGGCTTAATCTCCGC-3’

Fw-GAPDH: 5’-ACCACAGTCCATGCCATCAC-3’

Rv-GAPDH: 5’-TCCACCACCCTGTTGCTGTA-3’

### In Vivo Analysis of Neuronal Activity

CaMPARI2 (Addgene: pAAV_hsyn_NES-his-CaMPARI2-F391W-L398V; Plasmid #119723) was used to analyze neuronal activity of live neurons in culture. This sensor undergoes photoconversion from green to red in the presence of calcium and 405 nm light^25^. Neuronal cultures were infected with adenoviral particles to express CaMPARI2 at DIV3, and neuronal activity was analyzed at DIV15. For the analysis, images (red-pre and green-pre) were first captured in an Axiovert200 inverted microscope (Zeiss) using a 20x objective in the red (561nm,) and green (488nm) channels. Then the field of view was illuminated for 5 sec with 405nm light. Then, images (red-post and green-post) were again captured in the red and green channels. Photoconversion levels were expressed for each cell using the following formula: (red-post/green-post) / (red-pre/green-pre). To correlate neuronal activity and Ng expression, glass-bottomed gridded plates were used, that make it easier to localize the same positions for in vivo imaging and post-fixation imaging. For neuronal activity, 9 different fields of view were captured as described. Cultured neurons were then fixed and processed for immunofluorescence with antibodies to Ng and MAP2. Finally, in vivo capture fields were localized and imaged to capture Ng and MAP2 staining.

### Analysis of HDAC4 Nucleus/Cytoplasm Localization

We analyzed the relative localization of endogenous HDAC4, or the expressed mutants HDAC4-Nuc and HDAC4-Cyto between the nucleus and cytoplasm. LSM710 confocal microscope (Zeiss) was used to capture DAPI and HDAC4 images, with a 40x oil immersion objective, capturing 25 fields per sample. In CellProfiler 3.0^26^, a pipeline was created where the DAPI images were used to segment cell nuclei, and HDAC4 images to define the plasma membrane contours. The cytoplasm was defined as the area between the plasma membrane and the nucleus. Then, the mean intensity of HDAC4 staining was measured in the nuclear and cytoplasmic masks of each cell and the ratios between them calculated.

### Bioinformatics

The human (hg38) and mouse (mm10) genomes were analyzed using the UCSC Genome Browser^27^. For visualizing sci-ATAC-seq in vivo and in vitro data reported by Sinnamon et al.^28^, paired-end FASTQ read files for all samples were downloaded from the Sequence Read Archive (SRA; runs: SRR7749424-SRR7749428). The read 1 files (forward reads) and read 2 files (reverse reads) were concatenated separately. Barcodes, present in the last 36 bp of the read 2 files, were separated using SeqKit (v. 2.1.0)^29^. The resulting read and barcode files were passed as input to scATAC-pro (v. 1.5.1)^30^ using the mouse mm10 genome reference and default parameters. The filtered alignment file from scATAC-pro, containing aligned reads for all the cells, had its read names trimmed to leave only the barcode sequence. This file was then split by cell type using the bam-split function from scitools (github.com/adeylab/scitools), with barcode and cell type annotation files^28^ from NCBI Gene Expression Omnibus (GEO), accession number GSE118987. The resulting files, containing mapped reads for each cell type, were visualized together using Integrative Genomics Viewer (IGV)^31^ with all the called peaks from the combined in vivo and in vitro analysis.

### Statistical Analysis

Statistical analysis was performed using GraphPad Prism 7 software. When comparing two experimental groups, the t-Student test was used. When comparing multiple experimental groups, a one-way ANOVA analysis with Bonferroni correction test was used. Data are expressed as mean ± standard error of the mean (SEM), and p values less than 0.05 were considered significant (*, p<0.05; **, p<0.01; ***, p<0.001).

## Results

### In vivo and in vitro Ng expression

In our previous study^19^, we identified that Ng expression in primary E18 rat hippocampal cultures begins in the second week and peaks in the third week, similar to other synaptic proteins. However, Ng expression was restricted to about 15% of excitatory neurons, contrasting with its higher and more widespread expression in adult hippocampal tissue. This led us to hypothesize that Ng expression is influenced by developmental events not reproduced in vitro, possibly related to the establishment of appropriate synaptic connectivity and activity or to more advanced stages of neuronal maturation. Our attempts to upregulate Ng expression in vitro, including the establishment of cultures from hippocampi obtained at E18, E20, P0, P2 or P5, showed no significant increases. We next examined Ng expression in organotypic cultures of hippocampal slices, a model that preserves tissue architecture. Using slices from P5-P6 rats we found that Ng decreased in the first week of culture and recovered to reach maximal levels from DIV11 (Figure 1A). Ng was widely present in the pyramidal and granular layers, but its levels were about 10 times lower than in adult tissue. We further explored Ng expression using the model of adult neurogenesis. Newborn dentate gyrus (DG) neurons, which recapitulate the maturation stages of development and integrate into existing circuits^32^, exhibited elevated levels of Ng expression comparable to those in preexisting neurons. To investigate the timing of Ng expression in nascent granule neurons, we analyzed its colocalization with various neurogenesis markers: doublecortin (DCX), calretinin (Calret), NeuN, and calbindin (Calbd) (Figure 1B). Immunofluorescence analysis revealed minimal overlap with early markers: 4.9% with DCX and 6.2% with Calret. In contrast, 96.5% of neurons expressed both NeuN and Ng. NeuN appears in the third week of postmitotic neuron life and is present in all granule neurons. Similarly, 91.6% of Ng-positive neurons expressed Calbd. These results suggest that Ng expression begins after NeuN but before Calbd, occurring between 4 and 8 weeks postnatal, during the final stages of neuronal maturation when newborn neurons integrate into preexisting circuits (Supp. Figure 1).

**Figure 1.**
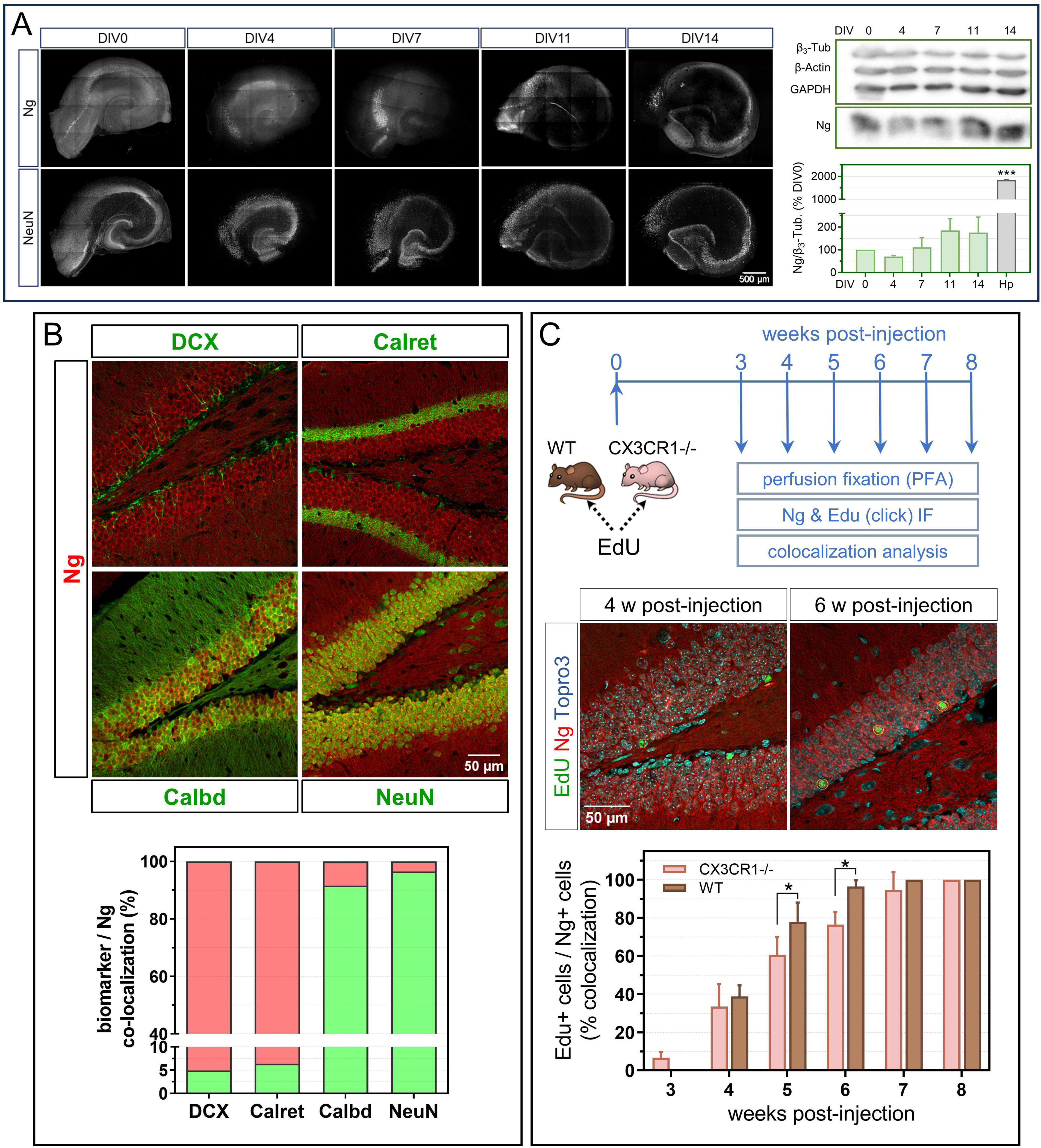
Ng expression in hippocampal organotypic cultures and newborn adult neurons. **A**. 400 µm slices of rat hippocampus (P5-P7) were maintained in culture and processed at different days in vitro (DIV) for western blot (WB) and immunofluorescence (IF) over two weeks. 5×5 image tiles of hippocampal slices incubated were acquired using anti-Ng and anti-NeuN antibodies (40X, Zeiss LSM710). Representative WB and quantification of Ng/β3Tubulin at different DIV and in intact adult tissue is shown (mean ± SEM, n=4). **B.** 2–3-month-old mice were anesthetized and perfused with 4% PFA/PBS. Their brains were cut into 50 µm thick sagittal sections and incubated with anti-Ng antibodies and antibodies against the neurogenesis markers DCX, Calretinin, NeuN and Calbindin to assess colocalization. Images acquired with a 20X objective (Zeiss LSM510). **C.** 2-month-old wild-type (wt) and CX3CR1 -/- mice were administered EdU intraperitoneally (IP) and sacrificed weekly from week 3 to week 8. Ng and EdU colocalization was analyzed in confocal images (Zeiss LSM510). Representative images of the dentate gyrus wt and CX3CR1 -/- mice 6 weeks after EdU administration are shown. The histogram shows the quantification of EdU/Ng colocalization at each time point (mean ± SEM, n=4).

We next used the cell cycle tracer 5-ethyl-2′-deoxyuridine (EdU) to label mitotic cells. Two-month-old mice were injected with EdU and sacrificed weekly, starting three weeks after injection and continuing for five weeks (Figure 1C). Three weeks after birth, EdU-labeled neurons were found in the DG subgranular layer (SGL), with no overlap between EdU and Ng expression. By the fourth week, 38.7% of EdU-labeled neurons expressed Ng, increasing to 96.4% at week six. By the seventh week, all EdU-labeled neurons expressed Ng. This indicates that Ng expression initiates between the fourth and sixth week after neuronal birth, coinciding with a period of intense synaptogenesis and the integration of newborn neurons into circuits^32,33^. These observations prompted us to investigate the effects of disrupting this integration process. We studied Ng expression in adult-born DG neurons from mice lacking the fractalkine receptor (CX3CR1 -/-), which is essential for neuron-microglia communication, and affects functional connectivity^34,35^ and synaptic maturation^36,37^. CX3CR1 -/- mice exhibit cognitive deficits and significant alterations in neuronal morphology and connectivity^38^. Comparing Ng expression in DG neurons from wild-type (wt) and CX3CR1 -/- mice after EdU labeling, we observed significant differences. CX3CR1 -/- mice exhibited reduced neurogenesis and delayed onset of Ng expression compared with wt mice (Figure 1C). These differences were most pronounced at six weeks after birth. However, by the eighth week, all EdU-labeled neurons from both groups expressed Ng, suggesting that neurons failing to integrate due to CX3CR1 deficiency may have been eliminated. These findings underscore the critical role of neuronal connectivity and synaptic activity in driving Ng expression.

### Synaptic activity regulates Ng expression

Spontaneous electrical activity in cultured hippocampal neurons begins by the end of the first week of culture and intensifies in subsequent weeks^39^, paralleling Ng expression. Previously, we demonstrated that long-term exposure to AP5, an NMDA receptor (NMDAR) antagonist, markedly reduces Ng expression, whereas exposure to NBQX, an AMPA receptor (AMPAR) antagonist, does not^19^. This prompted us to investigate whether direct stimulation with NMDA increases Ng expression. We found that only low doses (5 µM) and prolonged exposure to NMDA increased Ng levels (Figure 2A). Prolonged exposure to NMDA concentrations of 10 µM or higher induces neurotoxicity, as evidenced by observable morphological changes in neurons. Since AMPAR blockade does not affect Ng levels, NMDAR activity seems necessary to trigger the signaling cascade that activates Ng expression^4,40^.

**Figure 2.**
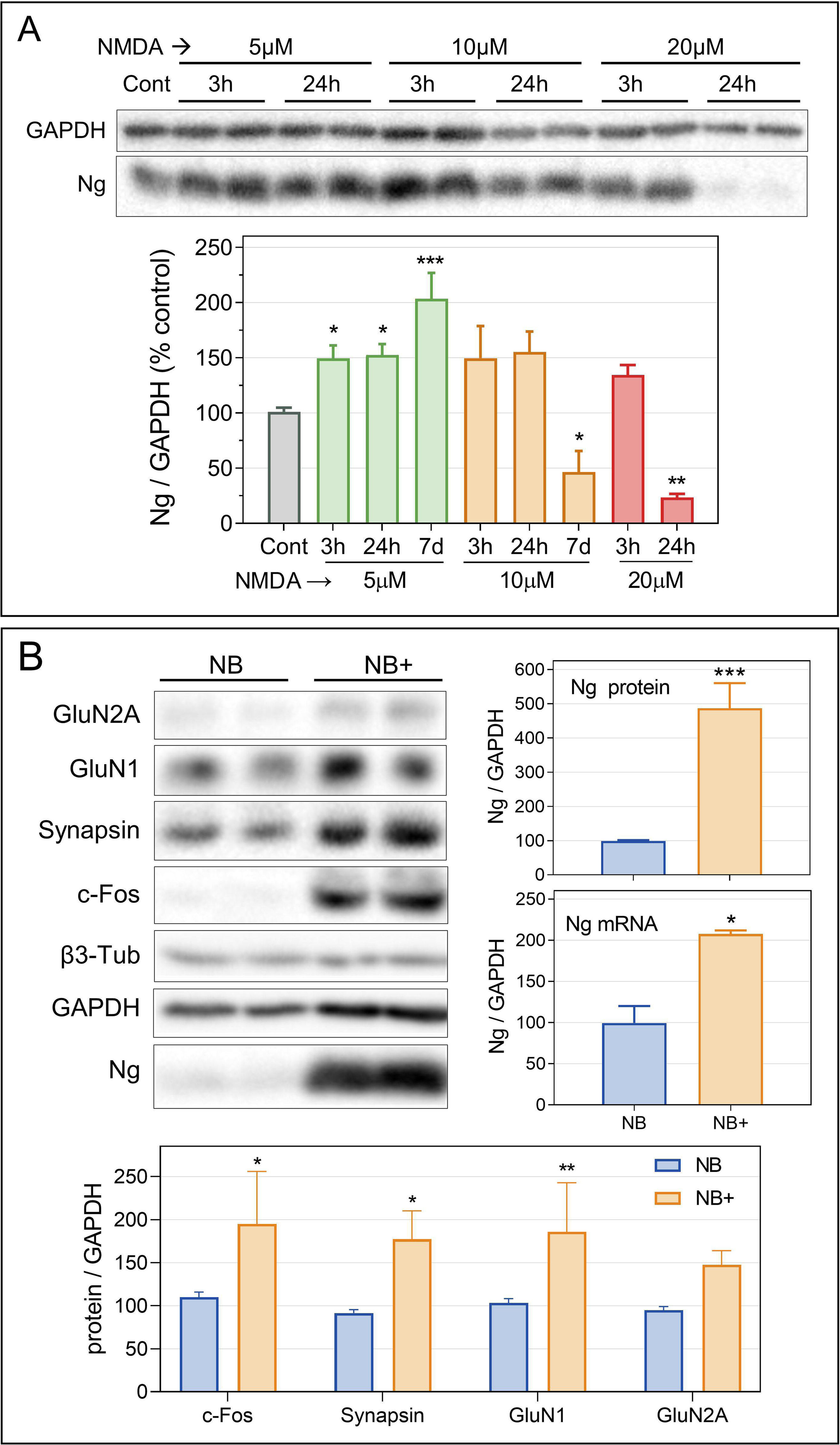
Increased Ng expression in synaptically active cultured neurons. **A.** DIV17 extracts of hippocampal neuron cultures were analyzed by WB following NMDA treatments (5-20 µM) for 3, 24 h or 7 days (mean ± SEM, n=3). **B.** Culture in NB+ medium boosts Ng expression. Hippocampal neuron cultures initially grown in NB medium were either maintained in NB or switched to NB+ medium from DIV7. Representative WB and quantification of Ng protein and mRNA levels are shown (mean ± SEM. Protein n=7; mRNA n=2). Protein levels of cFos, synapsin, GluN1 and GluN2A were also measured in DIV17 extracts from cultures grown in NB and NB+ (n=2).

We then compared the expression of Ng and other synaptic proteins in neurons cultured in Neurobasal^TM^ and B27^TM^ (NB) versus Neurobasal+™ and B27+™ (NB+), a newer formulation designed to enhance endogenous synaptic activity. NB+ cultures showed a nearly 5-fold increase in Ng protein and a 2-fold increase in Ng mRNA levels compared to NB (Figure 2B). Additionally, NB+ significantly elevated c-Fos, synapsin, and the GluN1 and GluN2A subunits of NMDARs. The percentage of Ng-expressing neurons in the cultures increased from less than 20% in NB to more than 50% in NB+ (Supp. Figure 2A). Neurons in NB+ also displayed more complex morphologies, characterized by greater dendritic extension and branching. Analysis of presynaptic bouton densities revealed a nearly 50% increase in both excitatory and inhibitory synapses in NB+ (Supp. Figure 2B), indicating enhanced connectivity and more complex synaptic networks.

To confirm increased neuronal activity in NB+ cultures, we employed the photoconvertible calcium sensor CaMPARI2^41^, which shifts fluorescence from green to red upon exposure to 405 nm light in the presence of calcium ions, indicating heightened activity. Neurons in NB+ exhibited significantly higher levels of CaMPARI2 photoconversion than those in standard NB (Supp Figure 3), confirming enhanced synaptic activity. Next, we correlated CaMPARI2 photoconversion with Ng levels, finding that neurons with low to medium Ng expression had low photoconversion, whereas those with medium to high Ng levels exhibited high photoconversion (Figure 3A). This suggests a relationship between Ng expression and synaptic activity. Additionally, we evaluated the expression of cFos, a marker of recent neuronal activity, and Ng in both NB and NB+ cultures using immunofluorescence. We observed over 90% colocalization between Ng and c-Fos in both NB and NB+ media, despite differences in the proportion of neurons expressing both proteins in each medium (Figure 3B). Again, these results suggest a close relationship between synaptic activity and Ng expression.

**Figure 3.**
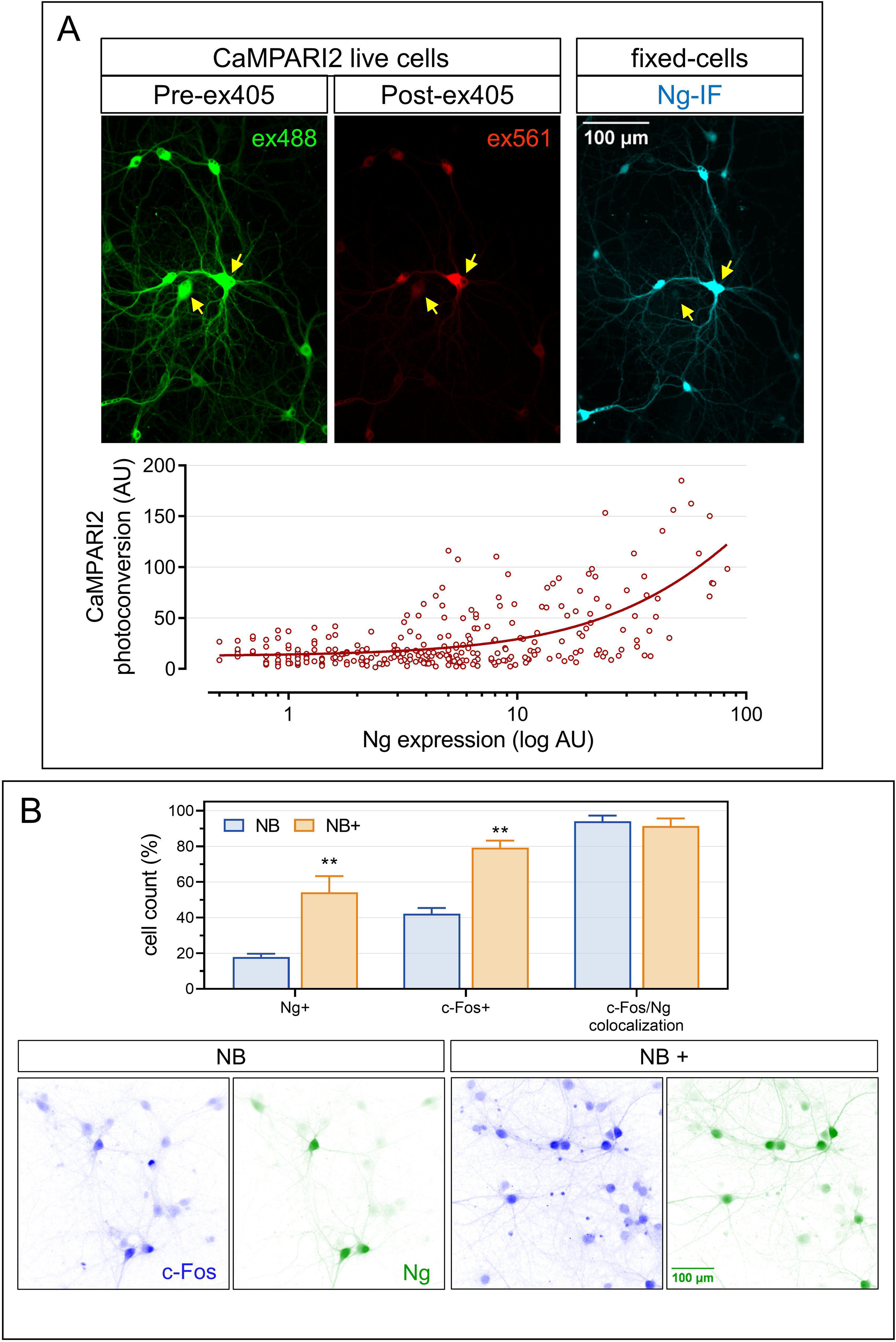
Synaptically-active neurons express higher Ng levels. **A.** Hippocampal neurons grown on glass-bottomed gridded plates were transduced at DIV3 with CaMPARI2 AAVs and analyzed for in vivo photoconversion at DIV15. Following photoconversion, neurons were processed for IF with anti-Ng antibodies. Ng content and CaMPARI2 photoconversion were measured in individual neurons. The image set highlights two neurons with similar CaMPARI2 expression (ex488, Pre-ex405) but markedly distinct levels of CaMPARI2 photoconversion and Ng. The graph below compares CaMPARI2 photoconversion and Ng levels in individual neurons, with the fitted exponential curve indicating a direct correlation between neuronal activity (CaMPARI2 photoconversion) and Ng expression. **B.** Hippocampal neurons cultured in NB and NB+ media were processed for IF with anti-cFos and anti-Ng antibodies at DIV16. The colocalization of both proteins was analyzed. Images acquired with a 25X objective (Zeiss Axiovert 200M). The upper histogram represents the percentage of cells expressing Ng, cFos and their colocalization in neurons cultured in NB and NB+ media (mean ± SEM, n=3). I_Blue and I_Forest lookup tables from https://github.com/cleterrier/ChrisLUTs were used.

### NMDAR activity drives Ng expression

To investigate the involvement of NMDARs in the increased Ng expression observed in NB+ medium, we administered various NMDAR antagonists at different time points (DIV7, DIV10, or DIV14) and maintained treatment until DIV17. MK-801, which selectively blocks active NMDARs, AP5, or Ifenprodil, a selective GluN2B antagonist, reduced Ng levels to those comparable to NB. In contrast, Memantine, a NMDAR antagonist with higher affinity for extrasynaptic NMDARs^42,43^, and TCN-201, a selective antagonist of GluN2A, did not decrease Ng expression (Figure 4A). These effects were more pronounced with longer treatments, suggesting that that NMDAR activity may modulate maturation processes that subsequently influence Ng expression.

**Figure 4.**
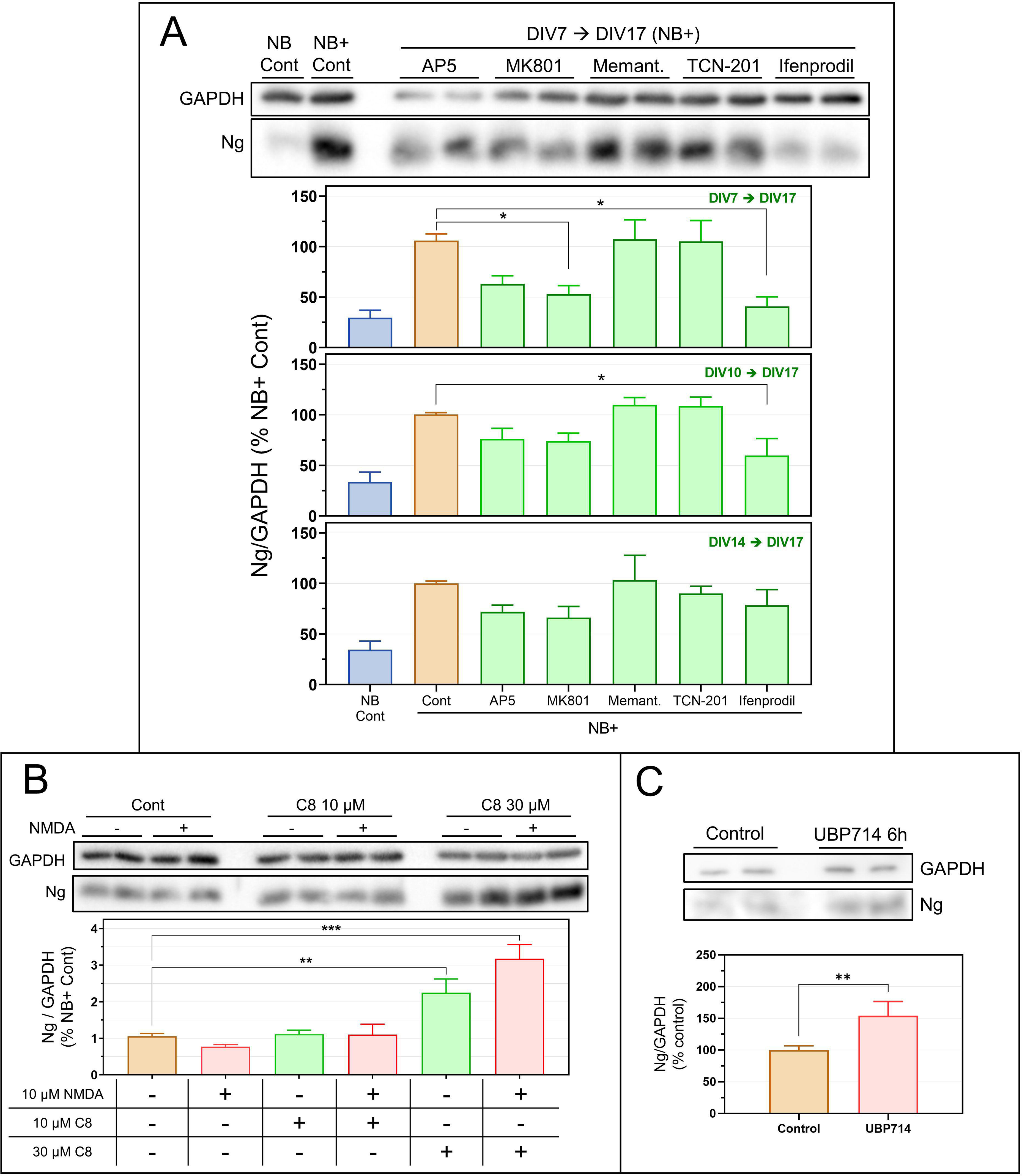
NMDAR activity drives Ng expression in cultured neurons. **A.** Hippocampal neuron cultures maintained in NB+ medium from DIV7 were treated with various NMDARs antagonists for 10, 7 or 3 days before extraction and analysis by WB at DIV17. A representative WB after 10-day treatments is shown. Treatments included AP5 (50 µM), MK801 (20 µM), Memantine (2 µM), TCN201 (10 µM) and Ifenprodil (10 µM). Differences in Ng expression were analyzed with respect to control neurons (NB+ Cont) without treatment (mean ± SEM, n=3). **B.** Hippocampal neuron cultures maintained in NB+ medium from DIV7 were treated with the C8 compound (10 or 30 µM) for 3 or 6 hours, in the presence or absence of NMDA (10µM) and processed for WB at DIV17 (mean ± SEM, n=3). **C.** Hippocampal neuron cultures maintained in NB medium were treated with the positive allosteric modulator (PAM) UBP714 (20 µM) for 6 hours and processed for WB at DIV16 (mean ± SEM, n=4).

NMDARs play distinct roles based on their localization: synaptic activation promotes neuroprotection and CREB-dependent transcription, whereas extrasynaptic activation induces apoptosis and suppresses synaptic gene transcription^44,45^. Given that Ng expression in NB+ depends on NMDAR activity, we investigated the impact of blocking extrasynaptic NMDARs. Yan et al.^46^ reported that extrasynaptic NMDAR toxicity involves interactions between GluN2A and GluN2B subunits with TRPM4 channels. Compound C8^46^ disrupts this interaction, preventing pro-apoptotic cascades without affecting neuroprotective signaling. In our study, application of compound C8 (30 μM) to neurons cultured in NB+ increased Ng levels, particularly in the presence of 10 μM NMDA (Figure 4B). This suggests that both synaptic and extrasynaptic NMDARs are active in NB+, with extrasynaptic NMDARs partially masking the effects of synaptic NMDARs. Another strategy used was the use of positive allosteric modulators (PAMs) of NMDARs. These compounds enhance the effectiveness of the endogenous natural agonist, synaptically released glutamate. Short-term treatments (6 hours) with the NMDAR PAM UBP714 resulted in a 50% increase in Ng levels in neurons cultured in NB medium (Figure 4C). These results further support the view that synaptic activation of NMDARs enhances Ng expression.

### HDAC4 suppresses the induction of Ng expression stimulated by NMDA

Histone deacetylase 4 (HDAC4), a class IIa histone deacetylase, plays a crucial role in activity-driven transcriptional regulation. Its shuttling between the nucleus and cytoplasm is regulated by NMDAR activity^47,48^. To investigate HDAC4’s role in Ng expression, we analyzed its subcellular localization in our neuronal cultures after 6 hours of AP5 or NMDA exposure. AP5 caused HDAC4 accumulation in the nucleus, particularly noticeable in neurons cultured in NB+ (Supp. Figure 4A). In contrast, NMDA reduced the nucleus-to-cytoplasm ratio more in NB than in NB+ medium. Additionally, NMDA intensified HDAC4 localization in dendrites, contrasting with its perinuclear localization in untreated neurons and those treated with AP5.

To study the impact of HDAC4 on Ng expression, we employed two HDAC4 mutants: HDAC4-Nuc (HDAC4-3SA^49^), which localizes to the nucleus, and HDAC4-Cyto (HDAC4-Δ1-217), which is exclusively cytosolic (Supp. Figure 4B). We analyzed the expression of Ng and synapsin, a known target of HDAC4^47^, in neurons expressing these mutants following silencing of endogenous HDAC4 (Supp. Figure 5 and Figure 5). HDAC4-Nuc consistently reduced both Ng and Synapsin levels across all conditions, with a more pronounced effect in NB+ neurons. In contrast, HDAC4-Cyto increased Ng levels specifically in NB neurons, suggesting that HDAC4 suppresses Ng expression in neurons cultured in NB medium. Silencing endogenous HDAC4 was essential to uncovering this effect. Given the high basal levels of Ng in NB+ neurons, it appears that increased synaptic activity counteracts HDAC4-mediated repression.

**Figure 5.**
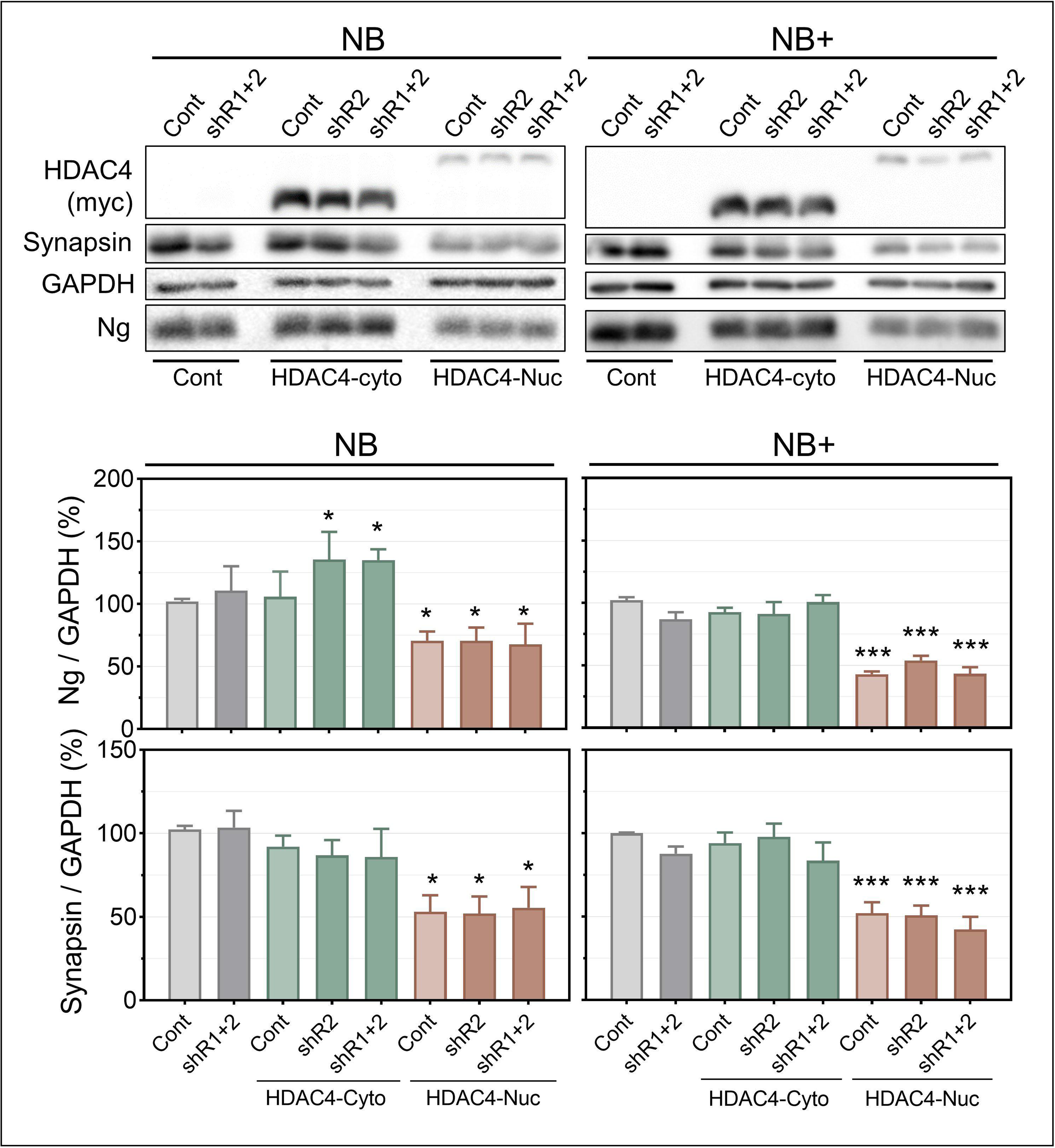
HDAC4-Nuc decreases Ng expression. Cultures of hippocampal neurons were transduced with HDAC4-Nuc or HDAC4-Cyto lentivirus at DIV9, with or without endogenous HDAC4 silencing from DIV15, and processed for WB at DIV17. Ng and synapsin levels were normalized by GAPDH and compared to their respective (non-transduced) controls in NB and NB+ media (mean ± SEM, n=3).

To further explore HDAC4’s repressive role in Ng expression, we treated hippocampal neurons expressing HDAC4-Nuc with NMDA (5μM) for either 24 hours or 7 days. Consistent with previous findings, prolonged low-level NMDA exposure significantly increased Ng expression. HDAC4-Nuc reduced Ng and synapsin levels in both NB and NB+ neurons and blocked the positive effect of NMDA (Figure 6A). These findings suggest that HDAC4 actively mediates the NMDA-induced increases in Ng expression, consistent with the proposal by Sando et al.^47^ that NMDAR activity confines HDAC4 to the cytoplasm, thereby preventing its suppressive activity on synaptic gene expression such as Ng. Next, we assessed the impact of various HDAC inhibitors on Ng expression by treating neurons for 24 or 48 hours during the third week of culture. Notably, class IIa HDAC specific inhibitors like LMK-235, markedly increased Ng levels, especially in neurons cultured in NB and after 48 hours of exposure (Figure 6B). Inhibitors of other HDACs had minimal effect, indicating HDAC4’s direct role in regulating Ng expression. We also measured Ng mRNA levels in neurons treated with LMK-235 or expressing HDAC4-Nuc. In both NB and NB+ media, inhibition of endogenous HDAC4 by LMK-235 increased Ng mRNA levels, while HDAC4-nuc reduced them. Notably, Ng mRNA levels were consistently higher in neurons cultured in NB+ compared to NB medium (Figure 6C).

**Figure 6.**
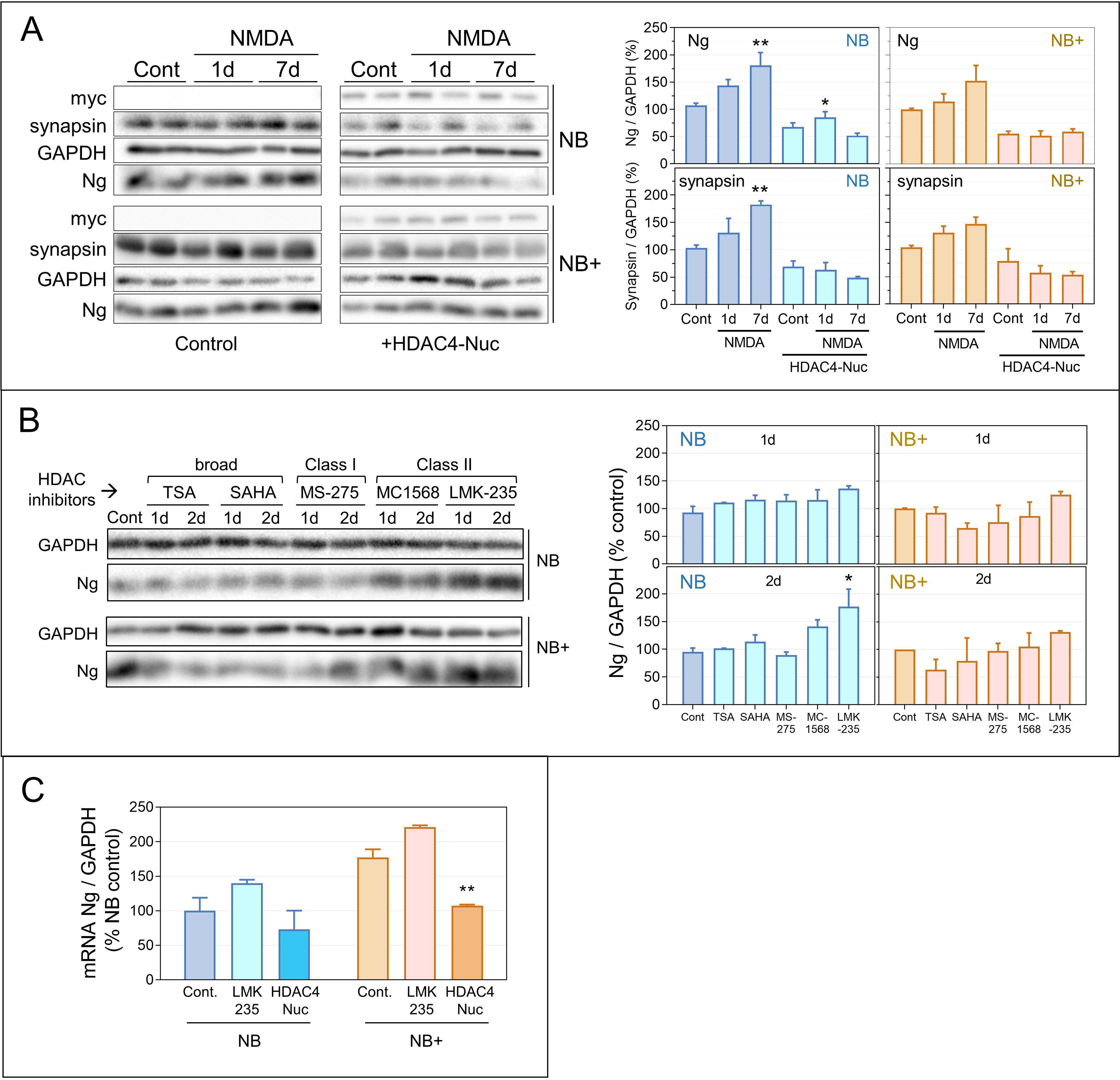
HDAC4 prevents NMDA-induced expression of Ng. **A.** Cultures of hippocampal neurons grown in NB or NB+ media were transduced with HDAC4-Nuc lentivirus at DIV9 and treated with 5 µM NMDA for 1 or 7 days. At DIV17, cultures were processed for WB using anti-Ng and anti-synapsin antibodies. Data in the histograms represent normalized levels corrected by GAPDH and referred to their respective non-transduced and non-treated controls. **B.** Cultures of hippocampal neurons were treated with the following HDAC inhibitors for 24 or 48 hours: TSA (250 nM), SAHA (1 µM), MS-275 (5 µM), MC1568 (5 µM) or LMK-235 (2 µM). Treatments were terminated at DIV16, and samples were processed for WB. Representative WBs are shown, and histograms present the quantification of normalized Ng levels in NB and NB+ media (mean ± SEM, n=2). **C.** RT-PCR of Ng mRNA. Cultures of hippocampal neurons maintained in NB or NB+ medium were transduced with HDAC4-Nuc lentivirus at DIV9 or treated with LMK-235 (2µM) for 48 hours. At DIV17, total RNA was prepared, and Ng mRNA levels were analyzed by RT-PCR (mean ± SEM, n=2). Differences were analyzed with respect to non-transduced, non-treated controls.

### NB+ enhances Ng gene transcription via NMDAR: Ng gene promoter analysis

Synaptic activity is well known to influence gene expression at the transcriptional level^4^. In this study, we investigated how NMDARs and HDAC4 regulate Ng expression in response to synaptic activity. To explore the mechanisms, we constructed a reporter using an AAV vector (pAAV-pNg2.4-mGL) containing a 2.4 Kbp sequence of the rat Ng gene promoter region driving the expression of the fluorescent protein mGreenLantern (mGL). Neurons were transduced with pAAV-pNg2.4-mGL virus at DIV3, maintained in NB or switched to NB+ at DIV7, and analyzed at DIV17 by Western blotting (WB) and immunofluorescence (IF) (Figure 7). Cultures in NB+ exhibited a significantly higher proportion of fluorescently labeled neurons (mGL) and elevated mGL levels measured by WB (Figure 7A, 7B), indicating enhanced Ng transcription driven by synaptic activity. To assess the role of NMDAR activity, NMDAR antagonists were added at DIV10 to neurons cultured in NB+ (Figure 7C). AP5, MK-801, and Ifenprodil significantly reduced mGL reporter levels, while Memantine caused a minor reduction and TCN-201 showed no effect. These results were mirrored in Ng expression, underscoring the involvement of GluN2B-containing NMDARs in mediating the effects of synaptic activity on Ng expression.

**Figure 7.**
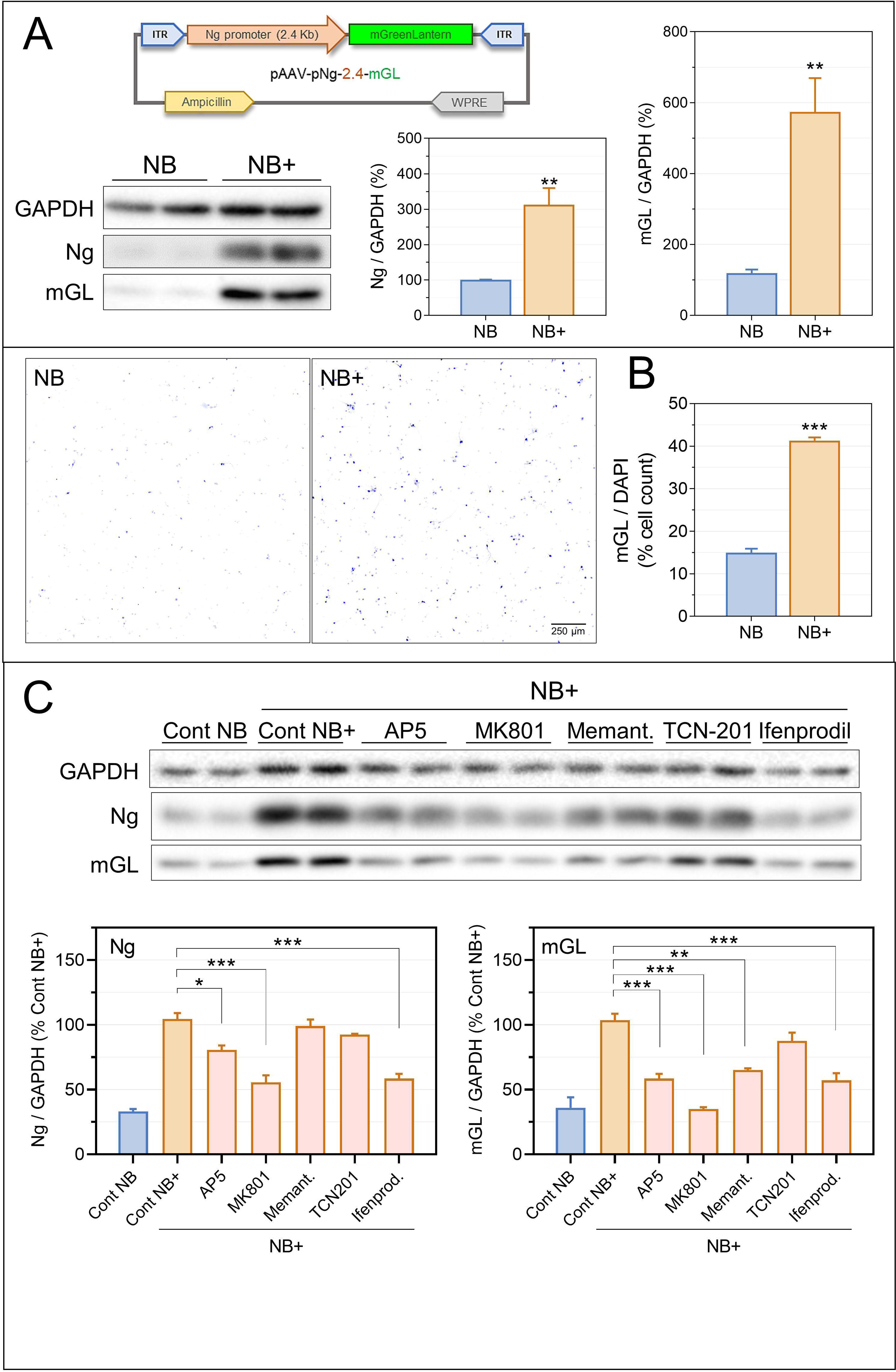
Analysis of Ng promoter reporter transcriptional activity. **A.** Schematic diagram of the pAAV-pNg2.4-mGL reporter. Ng and mGreenLantern (mGL) levels were measured at DIV17 in hippocampal neuron cultures maintained in NB or NB+ medium. Histograms depict the differences in Ng and mGL levels in neurons cultured in each medium (mean ± SEM, n=4). **B.** Representative images captured with a 20X objective (Zeiss Axiovert 200M) showing variations in the number of neurons expressing the mGL reporter in NB and NB+ medium. I_Blue lookup table from https://github.com/cleterrier/ChrisLUTs was used. **C.** Effect of various NMDAR antagonists on mGL reporter expression. Cultures of hippocampal neurons were transduced at DIV3 with pAAV-pNg2.4-mGL AAVs and switched to NB+ medium from DIV7. NMDARs antagonists were added at DIV10 and maintained up to DIV17. Representative WB and histograms illustrate mGL and Ng normalized levels (mean ± SEM, n=2). AP5 (50 µM), MK801 (20 µM), Memantine (2 µM), TCN201 (10 µM), Ifenprodil (10 µM).

To identify the most relevant promoter regions, we generated smaller fragments of the 2.4 Kbp sequence in the AAV constructs, each labeled according to its length (Figure 8A). Neurons transduced with these AAVs were cultured in NB or NB+, and mGL levels were analyzed by WB at DIV17. In NB medium, mGL expression was generally low, with neurons transduced with AAVs carrying the 2.0 or 1.4 Kbp fragments showing even lower levels compared to the 2.4 Kbp promoter. However, neurons with smaller fragments (0.85 and 0.23 Kbp) showed higher mGL expression. In NB+ medium, neurons with 2.0 and 1.4 Kbp fragments also had reduced expression, but the smaller fragments (0.85 and 0.23 Kbp) maintained similar levels to the 2.4 Kbp promoter. These findings suggest that the sequence between 2.4 and 2.0 Kbp contains elements that enhance transcription, as its deletion reduced reporter expression. On the other hand, the sequence between 2.0 and 0.85 Kbp contains elements that suppress transcription, as its absence increased expression. Therefore, neurons cultured in NB likely exhibit constitutive repression of Ng expression, which is alleviated by deleting the region between 2.4 to 0.85 Kbp. AAVs with the 0.85 and 0.23 Kbp promoter regions bypass this repression, achieving higher expression levels.

**Figure 8.**
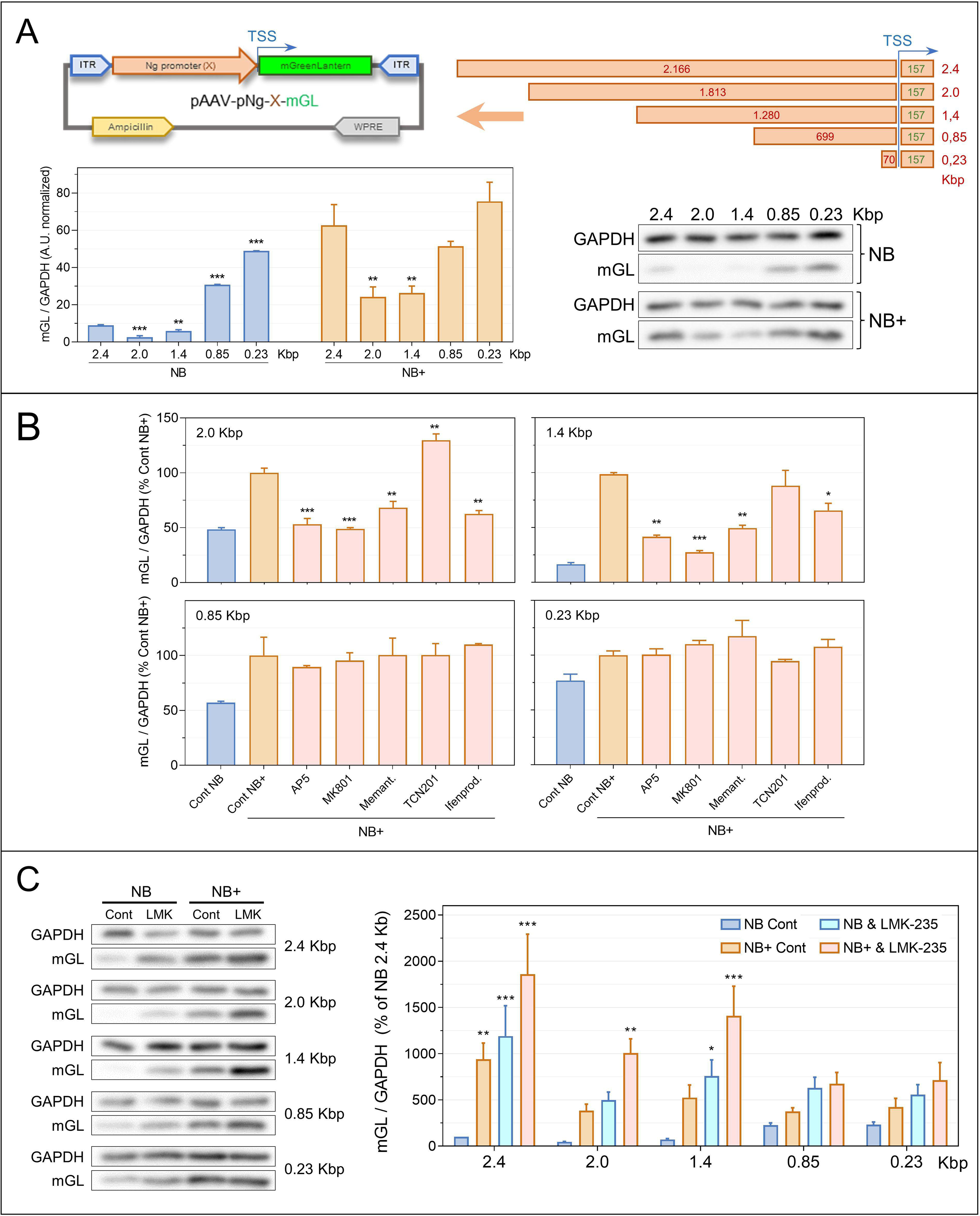
Analysis of Ng gene promoter fragments activity. **A.** Schematic diagram illustrating the different Ng promoter fragments utilized. Hippocampal neurons were transduced with AAVs carrying these promoter fragments at DIV3. Cultures were maintained in NB or switched to NB+ from DIV7. At DIV17, cell extracts were processed for WB, and the levels of mGL measured were normalized and presented as indicators of promoter activity (mean ± SEM, n=2). **B.** Effect of various NMDAR antagonists on promoter activity. Hippocampal neurons were transduced at DIV3 with AAVs carrying various Ng promoter activity reporters. From DIV7, cultures were maintained in NB+ medium. Starting from DIV10, NMDAR antagonists were added until DIV17. Promoter activity was assessed by quantifying mGL levels and analyzing differences compared to the NB+ control (mean ± SEM, n=2). The antagonists used were: AP5 (50 µM), MK801 (20 µM), Memantine (2 µM), TCN201 (10 µM), and Ifenprodil (10 µM). **C.** Analysis of HDAC4 inhibition on Ng promoter activity. Hippocampal neurons were transduced at DIV3 with AAVs carrying either the full 2.4 Kb Ng promoter (AAV-pNg 2.4-mGL) or fragments of 2.0, 1.4, 0.85, and 0.23 Kbp. Cultures were maintained in NB or NB+ medium. The HDAC4 inhibitor LMK-235 (2 µM) was added at DIV14 and maintained for 48 hours. Promoter activity was assessed by measuring mGL expression by WB. Data are presented as mean ± SEM, n=3.

Next, we investigated how NMDAR activity affects the activity of each promoter region. NMDAR antagonists AP5, MK801, Memantine, and Ifenprodil suppressed the NB+-induced increase in reporter expression for the longer promoters (2.4 Kbp in Figure 7C and 2.0 and 1.4 Kbp in Figure 8B). However, this suppression was not observed for the smaller 0.85 and 0.23 Kbp fragments, where mGL expression remained high despite NMDAR blockade. These results suggest that NMDAR-responsive elements are located between the fragments 0.85 and 2.4 Kbp. To identify the Ng promoter regions influenced by HDAC4, we examined the effect of LMK-235 on mGL expression driven by various promoter fragments. Significant differences were observed with the 2.4, 2.0, and 1.4 Kbp promoters when comparing LMK-235-treated neurons to their controls in both NB and NB+ media. In contrast, smaller fragments (0.85 and 0.23 Kbp) showed minimal changes (Figure 8C). These results again highlight the importance of the 0.85 to 2.4 Kbp region in modulating the sensitivity of Ng expression to synaptic activity.

Given HDAC4’s lack of deacetylase activity, its repressive role likely involves interactions with scaffold proteins and transcription factors^50^. To better understand the transcriptional regulation of Ng, we searched for transcription factor binding sites (TFBS) within the most conserved regions of the Ng gene promoter in 100 vertebrate species^51^ using the JASPAR2024^52^ TFBS database (Supp. Figure 6A). We identified three small (130-200 bp) conserved regions between position −2.4 and −1.4 Kbp featuring binding sites for REST, RFX4 and KLF transcription factors, among others. It has been described that REST (Repressor Element 1 Silencing Transcription factor) recruits complexes of CoREST and HDACs proteins^53^ to its TFBS to inhibit the expression of genes involved in neuron plasticity^54^. Near the Ng gene transcription start site (TSS) within the 0.23 Kbp fragment, we identified TFBS for EGR1, SP1-9 and ELK1, strongly indicating regulation by synaptic activity. In addition, beyond the 2.4 Kbp sequence, approximately −3.5 Kbp from the Ng gene TSS, there is another highly conserved area with binding sites for the myocyte enhancer factor-2 (MEF2) family of transcription factors. MEF2 regulates an intricate transcriptional program in neurons that controls synapse development^55^. The interaction between HDAC4 and the MEF2, known for its repressive role^56^, suggests that HDAC4 may regulate Ng expression by influencing MEF2C activity.

To evaluate the importance of identified promoter regions in Ng transcription, we sought to assess the chromatin accessibility of these areas. Utilizing data (GEO accession number GSE118987) from a study by Dr. A.C. Adey’s team (OHSU, Portland, USA)^28^, which investigated the chromatin landscape of the mouse hippocampus with single-cell resolution using sci-ATAC-seq, we observed a substantial overlap between highly conserved regions and those with the greatest chromatin accessibility (Supp. Figure 6B). The chromatin accessibility of these regions was notably heightened in excitatory neurons in vivo, in good agreement with the expression patterns of Ng, which is absent in hippocampal interneurons. However, cultured hippocampal excitatory neurons (VT1) exhibited slightly reduced accessibility, particularly in more distal conserved regions that contain MEF2 binding sites. This diminished accessibility may contribute to the observed downregulation of Ng expression in cultured neurons compared to adult hippocampal tissue. Overall, these findings reinforce the view that nuclear HDAC4 could suppress Ng expression by interacting with transcriptional complexes involving REST or MEF2 transcription factors.

## Discussion

The central nervous system (CNS) is highly complex and vulnerable to disturbances that can significantly affect cognitive functions. With increasing life expectancy and the growing prevalence of neurodegenerative diseases, safeguarding cognitive abilities has become a primary goal for health systems. This study focuses on Ng, a protein essential for cognitive function. Because of its beneficial effect on neuronal health, understanding the factors that regulate Ng expression could open new strategies to help combat cognitive decline and maintain mental health. Our findings clearly demonstrate that synaptic activity, particularly the activation of GluN2B-containing NMDARs, plays a pivotal role in regulating Ng expression dynamics within hippocampal neurons. The interplay between NMDAR signaling and HDAC4 reveals a finely tuned mechanism by which neuronal activity influences Ng expression. Through a comprehensive analysis of the Ng gene promoter, we have identified key regulatory sequences that are crucial for the modulation of Ng expression by synaptic activity and HDAC4.

### Ng expression parallels the onset of synaptic activity in neuronal networks

First, we demonstrated that Ng expression coincides with key developmental stages, such as synaptogenesis and the integration of adult newborn neurons into existing circuits of the hippocampal DG. Using EdU labeling, we observed that Ng expression begins during a period of intense formation of synapses and neuronal connectivity, underscoring its role in neuronal maturation. Comparing wild-type and CX3CR1 -/- mice, we found that Ng expression is delayed by approximately one week in CX3CR1 -/- mice, where the integration of newborn adult neurons into preexisting hippocampal circuits is impaired^57,58^. This again highlights the close interrelationship between synaptic activity and Ng expression, as neuron-microglia communication is essential for synaptic remodeling and circuit maturation. Our in vitro studies further strengthened the link between synaptic activity and Ng expression. Neurons cultured in NB+ medium, which enhances synaptic activity and promotes more complex neuronal morphology and connectivity, showed much higher levels of Ng and other synaptic proteins (Figure 2B). Additionally, our live-cell imaging observations using the CaMPARI-2 reporter revealed that Ng-expressing neurons displayed a substantial level of synaptic activity Ng (Figure 3). All these results indicate that at least part of Ng expression is dependent on synaptic activity.

### Activation of NMDA receptors containing the GluN2B subunit promotes Ng expression

Exploring the molecular mechanisms regulating Ng expression, we found that NMDAR activation is crucial. Blocking NMDARs with specific antagonists notably reduced Ng levels (Figure 4A). NMDAR involvement was further supported by experiments using i) direct stimulation with NMDA (Figure 2A), ii) the compound C8 to prevent extrasynaptic NMDAR activation, and iii) NMDAR-specific PAMs to enhance endogenous agonist action. Unraveling the stimulatory effect of direct NMDA exposure was challenging. Our results indicated a bell-shaped response of Ng expression to NMDA: low concentrations (<1 µM) do not alter Ng, whereas concentrations of 10 µM and higher not only failed to induce Ng expression but also caused morphological stress and apoptosis. This aligns with previous studies^59^, reflecting the opposing actions of synaptic (pro-survival) and extrasynaptic (pro-apoptotic) NMDARs, since bath application of NMDA activates both types of receptors^44^. Furthermore, “compound 8” (C8), which interferes with the interaction between TPRM4 and extrasynaptic NMDARs^46^, led to more than a twofold increase in Ng levels, even in the absence of NMDA (Figure 4B). In addition, positive allosteric modulators (PAMs) of NMDARs, like UBP714, significantly elevated Ng levels (Figure 4C). PAMs potentiate NMDAR responses to endogenously released glutamate^60^. Taken together, all these results demonstrate that modulating NMDAR activity induces substantial changes in Ng levels.

The activation of NMDARs, recognized as master regulators of neurodevelopment and synaptic plasticity, leads to both ionotropic Ca2+ influx-dependent and non-ionotropic^61,62^ signaling events. These events link synapse activation to cascades that control synaptic strength, spine size, neuronal excitability and survival^63,64^. We found that GluN2B-containing NMDARs play a crucial role in stimulating Ng expression. Multiple signaling pathways could mediate this effect. For example, CaMKII phosphorylates^65^ and interacts^66,67^ with the GluN2B NMDAR subunit to enhance synaptic transmission^66^. Recent studies have shown that CaMKII’s role in LTP induction is structural, requiring the binding of phospho-T286-CaMKIIα to the GluN2B subunit, independent of its kinase activity toward other substrates^66,68^. Nevertheless, Ca2+/CaM-activated CaMKII phosphorylates multiple proteins, including the neuron-abundant histone deacetylase HDAC4.

### HDAC4 represses NMDAR-stimulated Ng expression

Class II histone deacetylases, such as HDAC4, directly bind to and repress myogenic transcription factors of the myocyte enhancer factor-2 (MEF-2) family^56^. This repression disappears when HDAC4 is phosphorylated by CaM kinases, creating a docking site for 14-3-3 proteins, and unmasking a nuclear export sequence (NES) in the C-terminus of HDAC4, leading to its export to the cytoplasm. In neurons, this shuttling between the nucleus and the cytoplasm depends on synaptic activity^47,69^. In the present study, we discovered a precise mechanism by which HDAC4 represses NMDAR-stimulated Ng expression. A mutant of HDAC4 (HDAC4-Nuc), which is constitutively localized in the nucleus, decreased normal levels of Ng and synapsin, a confirmed target of HDAC4, as well as the stimulatory effect of NMDAR activation (Figure 5-6A). This repression was notably stronger in neurons cultured in NB+ medium, which exhibit higher synaptic activity. Using specific inhibitors, we determined that endogenous HDAC4 consistently suppresses Ng expression, with a more pronounced effect observed in NB medium, characterized by reduced endogenous synaptic activity (Figure 6B). These findings suggest that during periods of heightened neuronal activity, synaptic NMDAR activity retains HDAC4 in the cytoplasm, thereby preventing it from suppressing Ng expression. In contrast, neurons cultured in NB medium, with lower synaptic activity, undergo more robust and sustained HDAC4-mediated repression. This was evident as Ng expression increased upon expression of the cytoplasm-localized form of HDAC4 (HDAC4-cyto) and silencing endogenous HDAC4, an effect not observed in neurons cultured in NB+.

We observed that activity-dependent Ng expression in response to external treatments develops gradually. The stimulatory effect of low NMDA becomes more pronounced over extended periods (Figure 2). Switching to NB+ medium at DIV7 results in significantly higher Ng expression at later stages (DIV16-18), while switching during the third week shows a reduced effect (data not shown). Experiments using NMDAR antagonists demonstrate that Ng expression inhibition is more effective over longer periods (DIV7-DIV17) than shorter periods (DIV14-DIV17) (Figure 4). Given HDAC4’s role in regulating Ng expression, we suggest that increased Ng expression due to heightened synaptic activity may stem from epigenetic changes, likely in the Ng gene promoter region. Supporting this idea is the sci-ATACseq data analysis that we conducted on data from Sinnamon et al.^28^, that used samples from adult hippocampal tissue and cultured hippocampal neurons. This analysis shows that interneurons (VT2 and INT), which do not typically express Ng, presented reduced chromatin accessibility in the most conserved regions of the Ng promoter (Supp. Figure 6). Moreover, excitatory neurons cultured in vitro (VT1) exhibit notably lower chromatin accessibility in these regions compared to their counterparts in vivo (GRN and NR2). This observation could explain the low percentage of excitatory neurons showing Ng expression when cultured in NB medium. It is plausible that synaptic activity inducing these epigenetic modifications and changes in Ng expression must coincide with periods of neuronal maturation marked by intense synaptogenesis and the establishment of connectivity that ultimately shapes neural network functionality. Supporting this perspective, Han et al.^70^ described Ng’s crucial role in the conversion of silent synapses to AMPAR-positive synapses in the mouse visual cortex during the critical period. These processes depend on both synaptic activity (visual experience) and Ng expression for their proper functionality.

### Identifying regulatory elements in the Ng gene promoter

In a previous study^7^, we demonstrated that Ng expression in the developing rat forebrain occurs in two waves: the first around birth and the second, much more extensive, in the second week of life. This last wave coincides with the translocation of Ng into distal dendrites and a crucial period of experience-dependent synaptic refinement, essential for optimizing neural circuit function^70^. To date, few studies have explored the mechanisms of Ng expression, despite its beneficial effects on synaptic stability and cognitive function^71–73^. Hypothyroidism is known to decrease Ng expression.^74,75^. A thyroid hormone receptor (TR)/retinoid X receptor (RXR) response element has been identified within the first intron of the Ng gene^76^. However, not all brain regions respond equally to thyroid hormones. Even within the cerebral cortex, neurons in the same cortical layer show different Ng expression. Therefore, additional mechanisms must be involved in regulating Ng expression.

To identify regions of the Ng gene promoter potentially involved in its transcriptional regulation, we generated a fluorescent reporter AAV construct, whose expression is driven by various-sized DNA fragments derived from the 5’ region of the rat Ng gene’s transcription start site (TSS). Using the longer promoter fragment (2.4 Kbp), we observed substantially higher reporter expression in neurons cultured in NB+ medium compared to neurons in NB medium. Additionally, blocking NMDARs with MK-801 and Ifenprodil significantly reduced reporter expression to the basal levels observed in NB medium. This inhibitory effect was especially pronounced in NB+ medium, indicating that increased synaptic and NMDAR activity drive the heightened promoter activity. These results suggest that the increased Ng expression observed in neurons cultured in NB+ medium may be regulated by DNA sequences present in the 2.4 Kbp fragment studied. When comparing the effects of different DNA fragments on reporter expression, we found that the distal regions (2.4 to 0.85 Kbp) contain elements that enhance expression, as their removal decreases reporter levels. Conversely, the proximal regions (0.85 and 0.23 Kbp) either lack repressor elements or contain positive elements, as the reporter’s expression remained similar to that observed with the longer promoter. Analyzing the effects of several NMDAR antagonists on reporter expression, we found no effect when using the shorter promoter regions (0.85 and 0.23 Kbp), suggesting that NMDAR-responsive elements are located in the distal region of the promoter. Similar results were observed when inhibiting endogenous HDAC4: while increases in Ng were evident with the larger promoters, these increases were considerably reduced or disappear with the shorter promoters (Figure 8). These results suggest the existence of regulatory elements in the 5’ region of the Ng gene, located between −2.4 and −0.85 Kbp of the TSS, where both the stimulatory effect of NMDAR activity and the inhibitory effect of nuclear HDAC4 overlap.

Based on these results, we explored potential transcription factors involved in regulating Ng gene transcription. With the assistance of the JASPAR 2024 database^52^, we identified DNA binding sites of transcription factors within the most conserved regions of the Ng gene promoter, through comparative analysis across 100 vertebrate species^27^. Our search revealed multiple binding sites of transcription factors regulated by synaptic activity in the most distal part of the studied promoter region (Supp Figure 7). This finding aligns with our observations indicating a heightened sensitivity of these regions to NMDAR inhibition. Additionally, we pinpointed a highly conserved region at approximately 3,500 bp upstream of the Ng gene’s transcription start site (TSS) as a prominent target of the transcription factor MEF2, whose activity is repressed by HDAC4^56^.

Here, we have investigated the regulation of Ng expression, a crucial protein for synaptic plasticity and cognitive function. Our findings underscore the important role of synaptic activity, specifically mediated by GluN2B-containing NMDARs, in modulating Ng expression. We identified HDAC4 as a relevant mediator between NMDAR signaling and Ng expression dynamics. In our experimental approach to analyze Ng gene promoter activity we identified key regulatory sequences crucial for modulating Ng expression in response to synaptic activity and HDAC4 activity. Our study contributes valuable insights into the mechanisms governing Ng expression, which are crucial for understanding the mechanisms underlying cognitive function and exploring potential therapeutic avenues in neurodegenerative disorders. Further investigation into these mechanisms promises to enhance our understanding of synaptic plasticity and the maintenance of cognitive health.

## Author Contributions

R.A. and E.M-B carried out the experiments, analyzed the data and participated in discussions. F.J.D-G. conceived the study, contributed to data analysis and wrote the manuscript. All authors have read and approved the final version of the manuscript.

## Acknowledgements

We thank Dr. Félix Hernández (CBM) for providing fractalkine receptor deficient (CX3CR1 -/-) mice, and Dr. Hilmar Bading (IZN, Heidelberg University) for providing the C8 and C18 compounds. We thank the Advanced Light Microscopy Core Facility (SMOA) and the Genomic Facility of the “Centro de Biología Molecular Severo Ochoa” (CBM) for assistance with the imaging studies and the analysis of the sciATACseq data. The authors R.A. (PEJ-2017-AI/BMD-6017, PEJD-2019-PRE/BMD-14947) and E.M-B. (PEJD-2018/PRE/BDM-8491) were supported by the “Comunidad de Madrid” and “Fundación Severo Ochoa”. This work was supported by the Spanish Ministry of Science, Innovation and Universities (grant RTI2018-098712-B-I00). We also thank “Fundación Ramón Areces” for providing institutional support to CBMSO.

---

**Supplemental Figure 1. Schematic of adult neurogenesis and specific markers in the dentate gyrus**. ML: Molecular layer. GCL: Granule Cell layer. SGL: Subgranular layer. Diagram created based on data from^77^.

**Supplemental Figure 2. Increased maturation and synaptic density in neurons cultured in NB+ medium. A.** Hippocampal neurons cultured in NB and NB+ were processed for IF with anti-Ng and anti-MAP2 antibodies at DIV17. Images acquired with a 25X objective (Zeiss Axiovert 200M). The histogram shows the percentage of Ng-positive neurons (mean ± SEM, n=6). I_Forest and I_Bordeaux lookup tables from https://github.com/cleterrier/ChrisLUTs were used. **B.** Culturing hippocampal neurons in NB+ medium results in higher synaptic density. Neurons grown on coverslips and maintained in NB or NB+ medium were processed for IF at DIV17, with anti-GAD6 (inhibitory), anti-VGluT (excitatory), and anti-MAP2 antibodies. Images were acquired with a 40X objective (Zeiss Axiovert 200M). Histograms depict the average number of puncta per neuron in each condition (mean ± SEM, n=3).

**Supplemental Figure 3. Increased CaMPARI2 photoconversion in neurons cultured in NB+ medium**. The schematic diagram depicts the CaMPARI2 sensor and its mechanism of action. Hippocampal neurons were transduced with CaMPARI2 AAVs at DIV3 and analyzed for CaMPARI2 photoconversion at DIV15. Live-cell images acquired before (Pre-ex405) and after (Post-ex405) 405 nm laser illumination (20X objective, Zeiss Axiovert 200M). Histogram shows the quantification of CaMPARI2 photoconversion ((Red/Green)post / (Red/Green)pre) in individual neurons cultured in NB and NB+ media (mean ± SEM; n=4).

**Supplemental Figure 4. Subcellular localization of endogenous HDAC4 and mutants HDAC4-Nuc and HDAC4-Cyto in cultured hippocampal neurons. A.** Localization of Endogenous HDAC4. Hippocampal neurons cultured in either Nb or NB+ media were treated with AP5 (50 µM) or AP5 + NMDA (10µM) for 6 h and then processed for IF with anti-HDAC4 antibodies. DAPI staining was used to define the nuclear area. Confocal images were acquired with a 40X objective (Zeiss, LSM710). Histogram represents the analysis of a minimum of 200 cells per condition in two independent experiments (mean ± SEM). **B.** Subcellular Localization of HDAC4 Mutants: Top: Schematic diagram of HDAC4 wild-type (wt) and the HDAC4-nuc and HDAC4-Cyto mutants, highlighting the major domains and serine residues phosphorylatable by CaMKII. Bottom: Hippocampal neurons cultured in NB or NB+ media were transduced at DIV9 with lentivirus to express HDAC4-Nuc or HDAC4-Cyto mutants. At DIV16, they were processed for IF with anti-HDAC4 antibodies (to detect endogenous HDAC4, histogram control) or anti-Myc antibodies (to detect the HDAC4 mutants) and counterstained with DAPI. Confocal images were acquired using a 25X objective (Zeiss LSM710). The mean labeling intensity in the nuclear and cytoplasmic region of each individual cell was calculated, divided, normalized to their respective controls in NB or NB+ media, and plotted in the histogram (left). A minimum of 230 cells were analyzed in two independent experiments (mean ± SEM). Representative images from cultures maintained in NB+ are shown (right). I_Blue and I_Forest lookup tables from https://github.com/cleterrier/ChrisLUTs were used.

**Supplemental Figure 5. Proof of the effectiveness of endogenous HDAC4 silencing.** Hippocampal neuron cultures maintained in NB medium were transduced at DIV4 with lentivirus carrying shRNA constructs (shRNA1 and shRNA2) targeting endogenous HDAC4. HDAC4 protein levels were analyzed by WB at DIV17. The histogram shows the normalized levels of HDAC4 (mean ± SEM, n=4), demonstrating the effectiveness of HDAC4 silencing by shRNA constructs.

**Supplemental Figure 6. Conserved regions of the Ng promoter and associated transcription factor binding sites. A.** A 6.5 Kbp region of human chromosome 11, including the first exon, part of the first intron, and a 4 Kbp upstream region of the Ng gene transcription start site (TSS), is shown with the most conserved regions among 100 vertebrate species (Cons 100 Verts). High (orange) and low (blue) conservation levels are highlighted. The expanded promoter region below indicates the major transcription factors with binding sites in these conserved regions and shows the coverage of the various promoter fragments used in this study. **B.** The panel displays a capture from the Integrative Genomics Viewer (IGV 2.17) illustrating chromatin accessibility in the Ng gene promoter region. Data were extracted from single cell ATACseq experiments^28^ performed on mouse hippocampal adult tissue (in vivo: GRN, INT and NR2) and cultures of hippocampal neurons (In vitro: VT1, VT2), that were uploaded to the NCBI Gene Expression Omnibus (GEO) database (reference GSE118987). The highlighted homologous regions shown in panel A are also shown in the presumptive Ng gene promoter region of the mouse (mm10) genome. Note that most conserved regions of the Ng promoter roughly overlap with areas of greater chromatin accessibility. Additionally, these chromatin regions show high chromatin accessibility in excitatory neurons (GRN, NR2 and VT1) compared to interneurons (INT and VT2). Cultured hippocampal neurons exhibit lower levels of chromatin accessibility in most conserved regions compared to their in vivo counterparts.

## References

1. Mateos-Aparicio, P. & Rodríguez-Moreno, A. Calcium Dynamics and Synaptic Plasticity. in Calcium Signaling (ed. Islam, Md. S.) 965–984 (Springer International Publishing, Cham, 2020). doi:10.1007/978-3-030-12457-1_38.

2. Hoeflich, K. P. & Ikura, M. Calmodulin in Action: Diversity in Target Recognition and Activation Mechanisms. Cell 108, 739–742 (2002).

3. Citri, A. & Malenka, R. C. Synaptic Plasticity: Multiple Forms, Functions, and Mechanisms. Neuropsychopharmacology 33, 18–41 (2008).

4. Yap, E.-L. & Greenberg, M. E. Activity-Regulated Transcription: Bridging the Gap between Neural Activity and Behavior. Neuron 100, 330–348 (2018).

5. Díez-Guerra, F. J. Neurogranin, a link between calcium/calmodulin and protein kinase C signaling in synaptic plasticity. IUBMB Life 62, 597–606 (2010).

6. Watson, J. B., Sutcliffe, J. G. & Fisher, R. S. Localization of the protein kinase C phosphorylation/calmodulin-binding substrate RC3 in dendritic spines of neostriatal neurons. Proc. Natl. Acad. Sci. 89, 8581–8585 (1992).

7. Alvarez-Bolado, G., Rodríguez-Sánchez, P., Tejero-Díez, P., Fairén, A. & Díez-Guerra, F. J. Neurogranin in the development of the rat telencephalon. Neuroscience 73, 565–580 (1996).

8. Domínguez-González, I., Vázquez-Cuesta, S. N., Algaba, A. & Díez-Guerra, F. J. Neurogranin binds to phosphatidic acid and associates to cellular membranes. Biochem. J. 404, 31–43 (2007).

9. Huang, K. P. et al. Neurogranin/RC3 enhances long-term potentiation and learning by promoting calcium-mediated signaling. J. Neurosci. 24, 10660–10669 (2004).

10. Hoffman, L., Chandrasekar, A., Wang, X., Putkey, J. A. & Waxham, M. N. Neurogranin Alters the Structure and Calcium Binding Properties of Calmodulin. J. Biol. Chem. 289, 14644–14655 (2014).

11. Petersen, A. & Gerges, N. Z. Neurogranin regulates CaM dynamics at dendritic spines. Sci. Rep. 5, 11135 (2015).

12. Saunders, T. et al. Neurogranin in Alzheimer’s disease and ageing: A human post-mortem study. Neurobiol. Dis. 177, 105991 (2023).

13. Kvartsberg, H. et al. Cerebrospinal fluid levels of the synaptic protein neurogranin correlates with cognitive decline in prodromal Alzheimer’s disease. Alzheimers Dement. 11, 1180–1190 (2015).

14. Xue, M. et al. Association of cerebrospinal fluid neurogranin levels with cognition and neurodegeneration in Alzheimer’s disease. Aging 12, 1–15 (2020).

15. Liu, W. et al. Neurogranin as a Cognitive Biomarker in Cerebrospinal Fluid and Blood Exosomes for Alzheimer’s Disease and Mild Cognitive Impairment. SSRN Electron. J. (2020) doi:10/gnf2qd.

16. Miyakawa, T. et al. Neurogranin null mutant mice display performance deficits on spatial learning tasks with anxiety related components. Hippocampus 11, 763–775 (2001).

17. Zhong, L. et al. Increased Prefrontal Cortex Neurogranin Enhances Plasticity and Extinction Learning. J. Neurosci. 35, 7503–7508 (2015).

18. Jeon, S. G. et al. Intrahippocampal injection of a lentiviral vector expressing neurogranin enhances cognitive function in 5XFAD mice. Exp. Mol. Med. 50, e461–e461 (2018).

19. Garrido-García, A. et al. Neurogranin Expression Is Regulated by Synaptic Activity and Promotes Synaptogenesis in Cultured Hippocampal Neurons. Mol. Neurobiol. 56, 7321–7337 (2019).

20. Pak, J. H. et al. Involvement of neurogranin in the modulation of calcium/calmodulin-dependent protein kinase II, synaptic plasticity, and spatial learning: A study with knockout mice. Proc. Natl. Acad. Sci. U. S. A. 97, 11232–11237 (2000).

21. Kaleka, K. S. & Gerges, N. Z. Neurogranin restores amyloid β-mediated synaptic transmission and long-term potentiation deficits. Exp. Neurol. 277, 115–123 (2016).

22. Kaech, S. & Banker, G. Culturing hippocampal neurons. Nat. Protoc. 1, 2406–2415 (2006).

23. Gascón, S., Paez-Gomez, J. A., Díaz-Guerra, M., Scheiffele, P. & Scholl, F. G. Dual-promoter lentiviral vectors for constitutive and regulated gene expression in neurons. J. Neurosci. Methods 168, 104–112 (2008).

24. Qiu, J. et al. Mitochondrial calcium uniporter Mcu controls excitotoxicity and is transcriptionally repressed by neuroprotective nuclear calcium signals. Nat. Commun. 4, 2034 (2013).

25. Trojanowski, N. F., Bottorff, J. & Turrigiano, G. G. Activity labeling in vivo using CaMPARI2 reveals intrinsic and synaptic differences between neurons with high and low firing rate set points. Neuron 109, 663–676.e5 (2021).

26. McQuin, C. et al. CellProfiler 3.0: Next-generation image processing for biology. PLOS Biol. 16, e2005970 (2018).

27. Nassar, L. R. et al. The UCSC Genome Browser database: 2023 update. Nucleic Acids Res. 51, D1188–D1195 (2023).

28. Sinnamon, J. R. et al. The accessible chromatin landscape of the murine hippocampus at single-cell resolution. Genome Res. 29, 857–869 (2019).

29. Shen, W., Le, S., Li, Y. & Hu, F. SeqKit: A Cross-Platform and Ultrafast Toolkit for FASTA/Q File Manipulation. PLOS ONE 11, e0163962 (2016).

30. Yu, W., Uzun, Y., Zhu, Q., Chen, C. & Tan, K. scATAC-pro: a comprehensive workbench for single-cell chromatin accessibility sequencing data. Genome Biol. 21, 94 (2020).

31. Thorvaldsdóttir, H., Robinson, J. T. & Mesirov, J. P. Integrative Genomics Viewer (IGV): high-performance genomics data visualization and exploration. Brief. Bioinform. 14, 178–192 (2013).

32. Gonçalves, J. T., Schafer, S. T. & Gage, F. H. Adult Neurogenesis in the Hippocampus: From Stem Cells to Behavior. Cell 167, 897–914 (2016).

33. Ge, S., Sailor, K. A., Ming, G. & Song, H. Synaptic integration and plasticity of new neurons in the adult hippocampus. J. Physiol. 586, 3759–3765 (2008).

34. Basilico, B. et al. Microglia shape presynaptic properties at developing glutamatergic synapses. Glia 67, 53–67 (2019).

35. Fleming, L. L. & McDermott, T. J. Cognitive Control and Neural Activity during Human Development: Evidence for Synaptic Pruning. J. Neurosci. 44, (2024).

36. Sheridan, G. K. & Murphy, K. J. Neuron–glia crosstalk in health and disease: fractalkine and CX3CR1 take centre stage. Open Biol. 3, 130181 (2013).

37. Bertot, C., Groc, L. & Avignone, E. Role of CX3CR1 Signaling on the Maturation of GABAergic Transmission and Neuronal Network Activity in the Neonate Hippocampus. Neuroscience 406, 186–201 (2019).

38. Bolós, M. et al. Absence of microglial CX3CR1 impairs the synaptic integration of adult-born hippocampal granule neurons. Brain. Behav. Immun. 68, 76–89 (2018).

39. Cohen, E., Ivenshitz, M., Amor-Baroukh, V., Greenberger, V. & Segal, M. Determinants of spontaneous activity in networks of cultured hippocampus. Brain Res. 1235, 21–30 (2008).

40. Benito, E. & Barco, A. The Neuronal Activity-Driven Transcriptome. Mol. Neurobiol. 51, 1071– 1088 (2015).

41. Moeyaert, B. et al. Improved methods for marking active neuron populations. Nat. Commun. 9, 4440 (2018).

42. Xia, P., Chen, H. S. V., Zhang, D. & Lipton, S. A. Memantine preferentially blocks extrasynaptic over synaptic NMDA receptor currents in hippocampal autapses. J. Neurosci. 30, 11246–11250 (2010).

43. Wu, Y.-N. & Johnson, S. W. Memantine selectively blocks extrasynaptic NMDA receptors in rat substantia nigra dopamine neurons. Brain Res. 1603, 1–7 (2015).

44. Hardingham, G. E. & Bading, H. Synaptic versus extrasynaptic NMDA receptor signalling: Implications for neurodegenerative disorders. Nat. Rev. Neurosci. 11, 682–696 (2010).

45. Hardingham, G. E. & Bading, H. Coupling of extrasynaptic NMDA receptors to a CREB shut-off pathway is developmentally regulated. Biochim. Biophys. Acta BBA - Proteins Proteomics 1600, 148–153 (2002).

46. Yan, J., Bengtson, C. P., Buchthal, B., Hagenston, A. M. & Bading, H. Coupling of NMDA receptors and TRPM4 guides discovery of unconventional neuroprotectants. Science 370, eaay3302 (2020).

47. Sando, R. et al. HDAC4 governs a transcriptional program essential for synaptic plasticity and memory. Cell 151, 821–834 (2012).

48. Mielcarek, M., Zielonka, D., Carnemolla, A., Marcinkowski, J. T. & Guidez, F. HDAC4 as a potential therapeutic target in neurodegenerative diseases: a summary of recent achievements. Front. Cell. Neurosci. 9, (2015).

49. Cohen, T. J. et al. The Histone Deacetylase HDAC4 Connects Neural Activity to Muscle Transcriptional Reprogramming*. J. Biol. Chem. 282, 33752–33759 (2007).

50. Fitzsimons, H. L. The Class IIa histone deacetylase HDAC4 and neuronal function: Nuclear nuisance and cytoplasmic stalwart? Neurobiol. Learn. Mem. 123, 149–158 (2015).

51. Blanchette, M. et al. Aligning Multiple Genomic Sequences With the Threaded Blockset Aligner. Genome Res. 14, 708–715 (2004).

52. Rauluseviciute, I. et al. JASPAR 2024: 20th anniversary of the open-access database of transcription factor binding profiles. Nucleic Acids Res. 52, D174–D182 (2024).

53. Maksour, S., Ooi, L. & Dottori, M. More than a Corepressor: The Role of CoREST Proteins in Neurodevelopment. eNeuro 7, (2020).

54. Ballas, N., Grunseich, C., Lu, D. D., Speh, J. C. & Mandel, G. REST and Its Corepressors Mediate Plasticity of Neuronal Gene Chromatin throughout Neurogenesis. Cell 121, 645–657 (2005).

55. Flavell, S. W. et al. Genome-Wide Analysis of MEF2 Transcriptional Program Reveals Synaptic Target Genes and Neuronal Activity-Dependent Polyadenylation Site Selection. Neuron 60, 1022–1038 (2008).

56. Miska, E. A. et al. HDAC4 deacetylase associates with and represses the MEF2 transcription factor. EMBO J. 18, 5099–5107 (1999).

57. Bolós, M. et al. Absence of microglial CX3CR1 impairs the synaptic integration of adult-born hippocampal granule neurons. Brain. Behav. Immun. 68, 76–89 (2018).

58. Faust, T. E., Gunner, G. & Schafer, D. P. Mechanisms governing activity-dependent synaptic pruning in the developing mammalian CNS. Nat. Rev. Neurosci. 22, 657–673 (2021).

59. McQueen, J. et al. Pro-death NMDA receptor signaling is promoted by the GluN2B C-terminus independently of Dapk1. eLife 6, e17161 (2017).

60. Burnell, E. S. et al. Positive and Negative Allosteric Modulators of *N*-Methyl-D-aspartate (NMDA) Receptors: Structure–Activity Relationships and Mechanisms of Action. J. Med. Chem. 62, 3–23 (2019).

61. Gray, J. A., Zito, K. & Hell, J. W. Non-ionotropic signaling by the NMDA receptor: Controversy and opportunity [version 1; referees: 2 approved]. F1000Research 5, 1–8 (2016).

62. Park, D. K., Stein, I. S. & Zito, K. Ion flux-independent NMDA receptor signaling. Neuropharmacology 210, 109019 (2022).

63. Tolias, K. F. et al. The Rac1-GEF Tiam1 Couples the NMDA Receptor to the Activity-Dependent Development of Dendritic Arbors and Spines. Neuron 45, 525–538 (2005).

64. Wyszynski, M. et al. Competitive binding of α-actinin and calmodulin to the NMDA receptor. Nature 385, 439–442 (1997).

65. Omkumar, R. V., Kiely, M. J., Rosenstein, A. J., Min, K.-T. & Kennedy, M. B. Identification of a Phosphorylation Site for Calcium/Calmodulindependent Protein Kinase II in the NR2B Subunit of the *N*-Methyl-D-aspartate Receptor*. J. Biol. Chem. 271, 31670–31678 (1996).

66. Bayer, K.-U., De Koninck, P., Leonard, A. S., Hell, J. W. & Schulman, H. Interaction with the NMDA receptor locks CaMKII in an active conformation. Nature 411, 801–805 (2001).

67. Bayer, K. U. et al. Transition from Reversible to Persistent Binding of CaMKII to Postsynaptic Sites and NR2B. J. Neurosci. 26, 1164–1174 (2006).

68. Tullis, J. E. et al. LTP induction by structural rather than enzymatic functions of CaMKII. Nature 1–8 (2023) doi:10.1038/s41586-023-06465-y.

69. Chen, Y., Wang, Y., Modrusan, Z., Sheng, M. & Kaminker, J. S. Regulation of neuronal gene expression and survival by basal NMDA receptor activity: A role for histone deacetylase 4. J. Neurosci. 34, 15327–15339 (2014).

70. Han, K.-S., Cooke, S. F. & Xu, W. Experience-Dependent Equilibration of AMPAR-Mediated Synaptic Transmission during the Critical Period. Cell Rep. 18, 892–904 (2017).

71. Stefansson, H. et al. Common variants conferring risk of schizophrenia. Nature 460, 744–7 (2009).

72. Ruano, D. et al. Association of the gene encoding neurogranin with schizophrenia in males. J. Psychiatr. Res. 42, 125–133 (2008).

73. Coldren, C. D. et al. Chromosomal microarray mapping suggests a role for BSX and Neurogranin in neurocognitive and behavioral defects in the 11q terminal deletion disorder (Jacobsen syndrome). Neurogenetics 10, 89–95 (2009).

74. Iñiguez, M. A., Rodriguez-Peña, A., Ibarrola, N., Morreale de Escobar, G. & Bernal, J. Adult rat brain is sensitive to thyroid hormone. Regulation of RC3/neurogranin mRNA. J. Clin. Invest. 90, 554–8 (1992).

75. Enderlin, V. et al. Retinoic acid reverses the PTU related decrease in neurogranin level in mice brain. J. Physiol. Biochem. 60, 191–198 (2004).

76. De Arrieta, C. M., Morte, B., Coloma, A. & Bernal, J. The human RC3 gene homolog, NRGN contains a thyroid hormone-responsive element located in the first intron. Endocrinology 140, 335–343 (1999).

77. Zhang, J. & Jiao, J. Molecular Biomarkers for Embryonic and Adult Neural Stem Cell and Neurogenesis. BioMed Res. Int. 2015, 727542 (2015).

