## Supplementary figures and images for "HDAC4 Inhibits NMDA Receptor-Mediated Stimulation of Neurogranin Expression"

### Supp. Figure 1

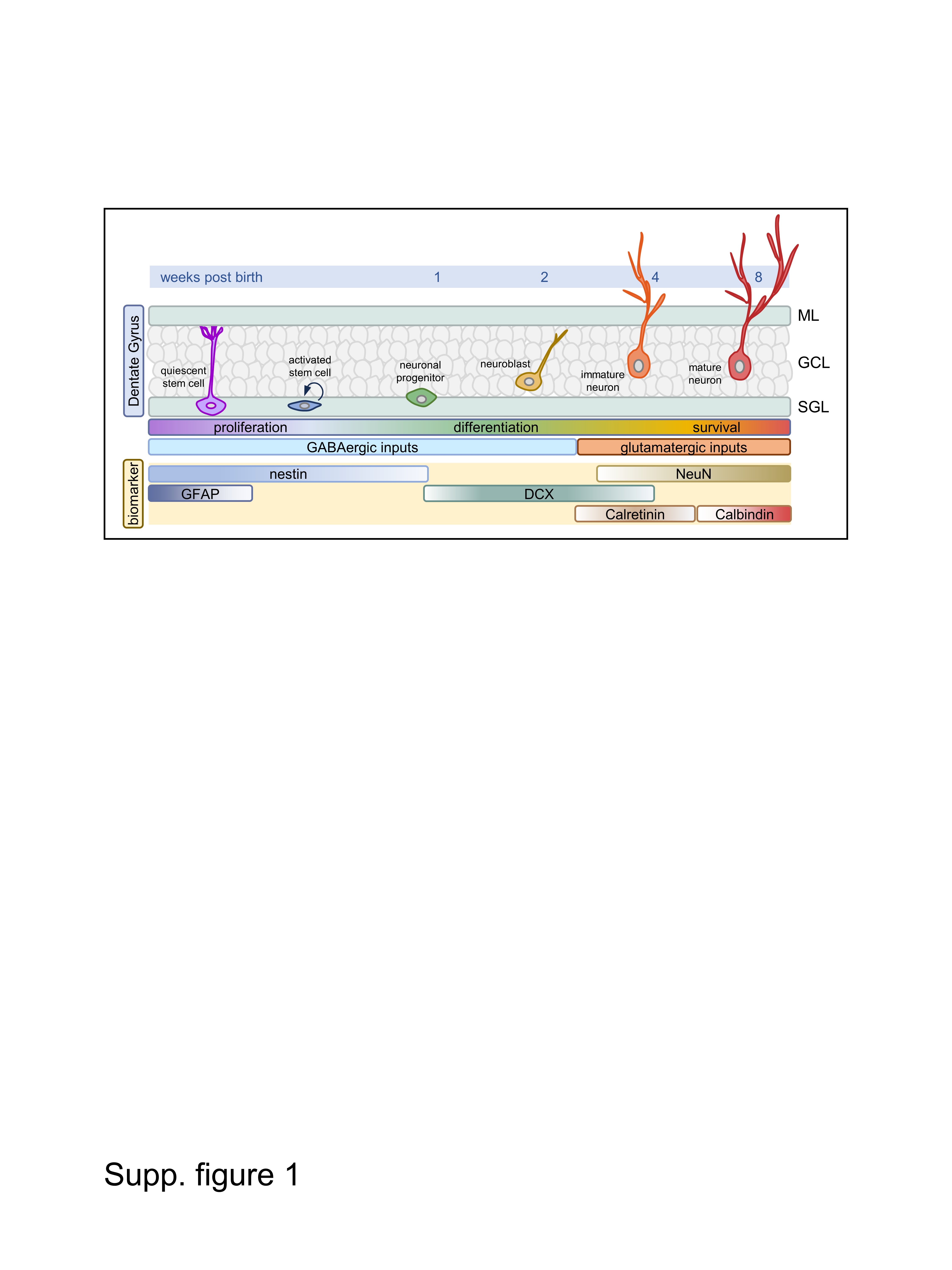

### Supp. Figure 2

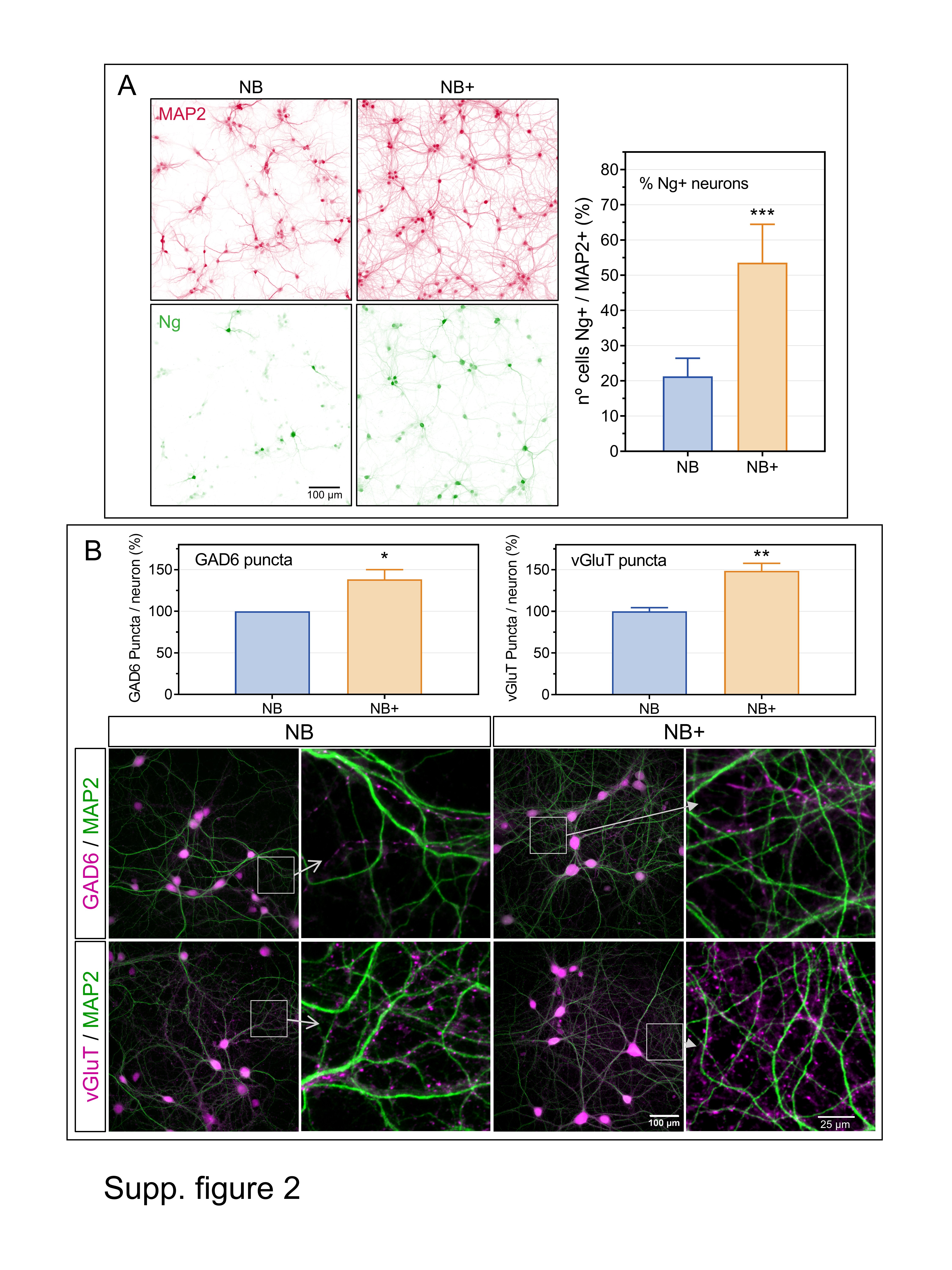

### Supp. Figure 3

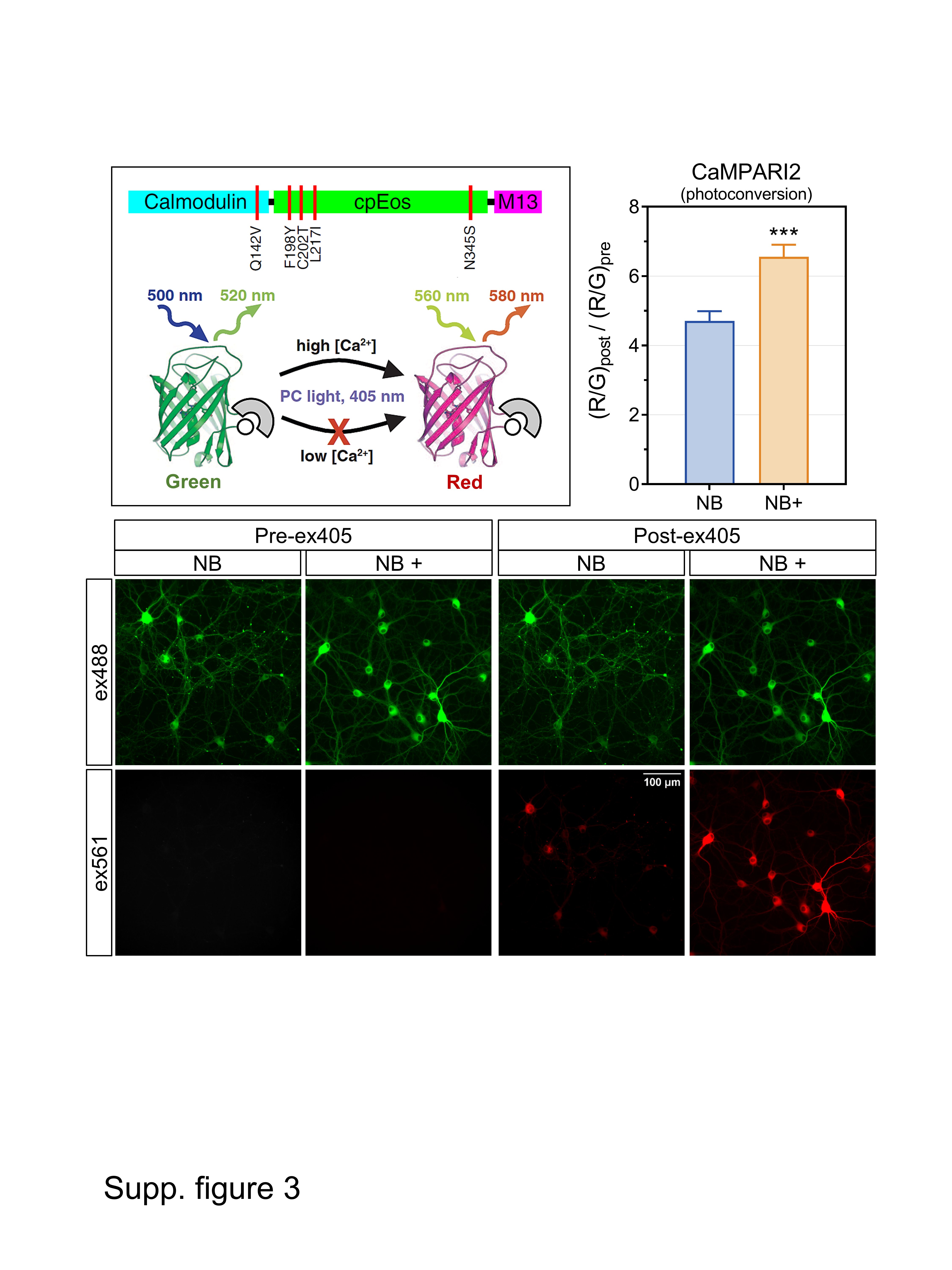

### Supp. Figure 4

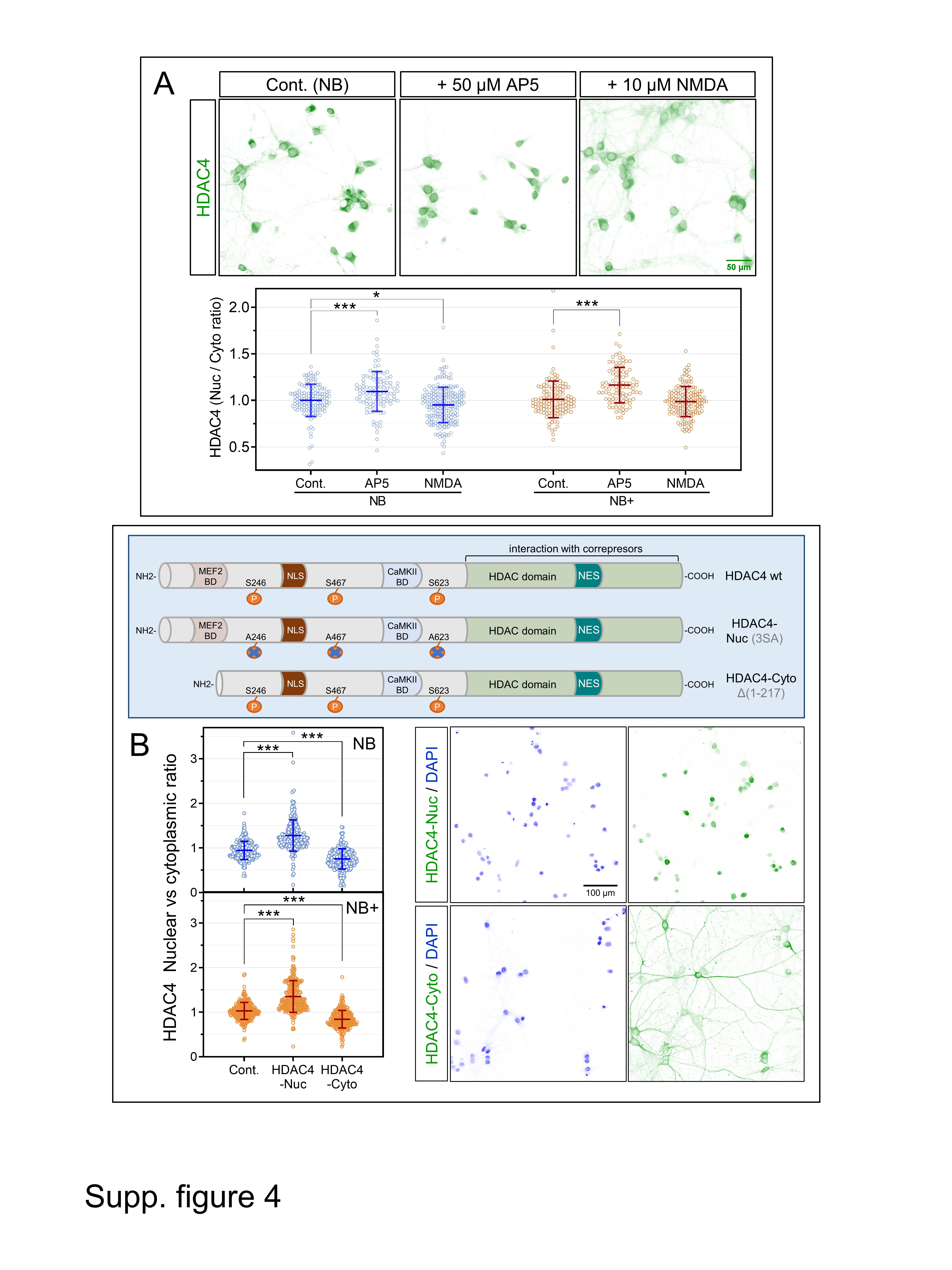

### Supp. Figure 5

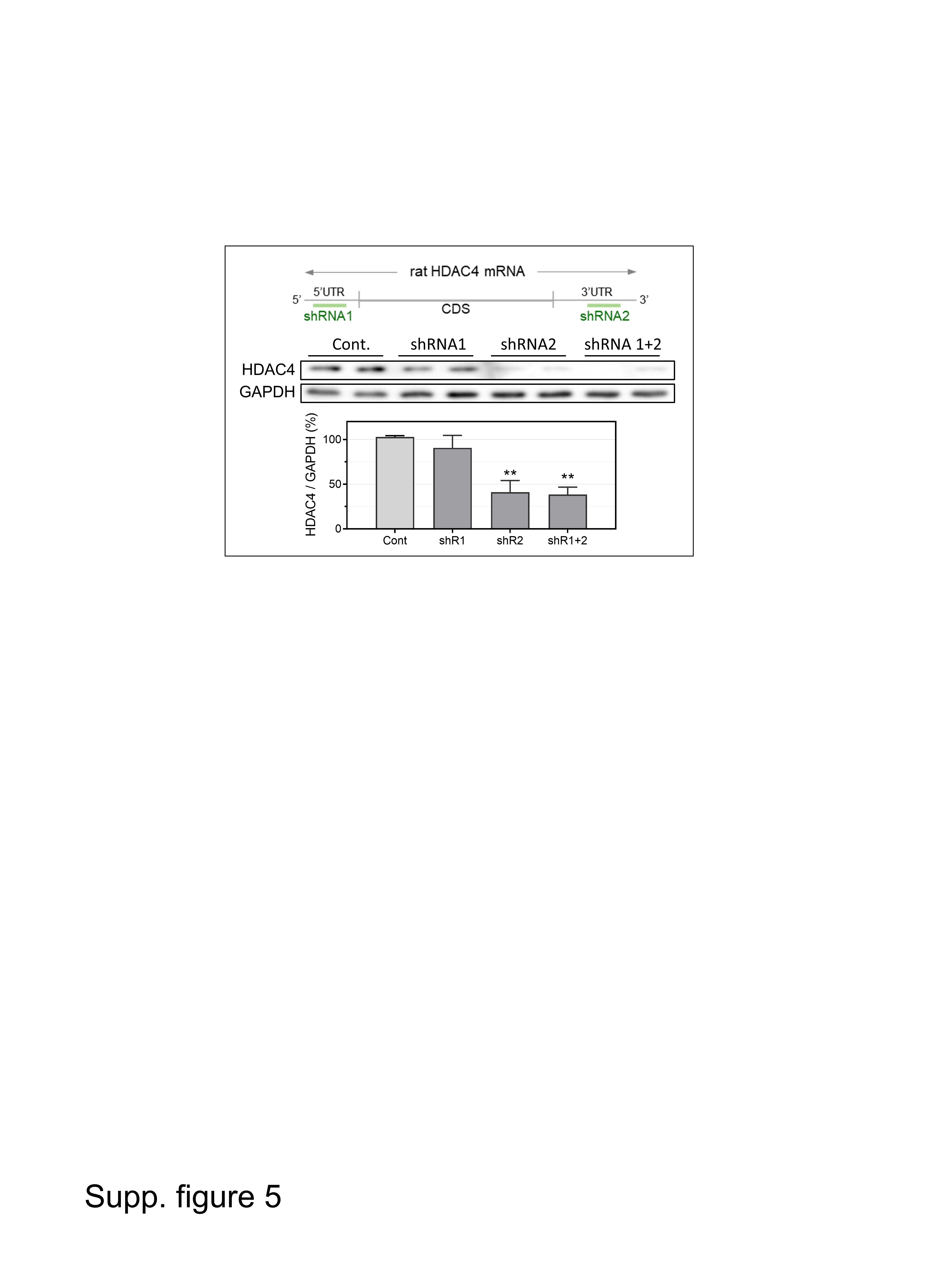

### Supp. Figure 6

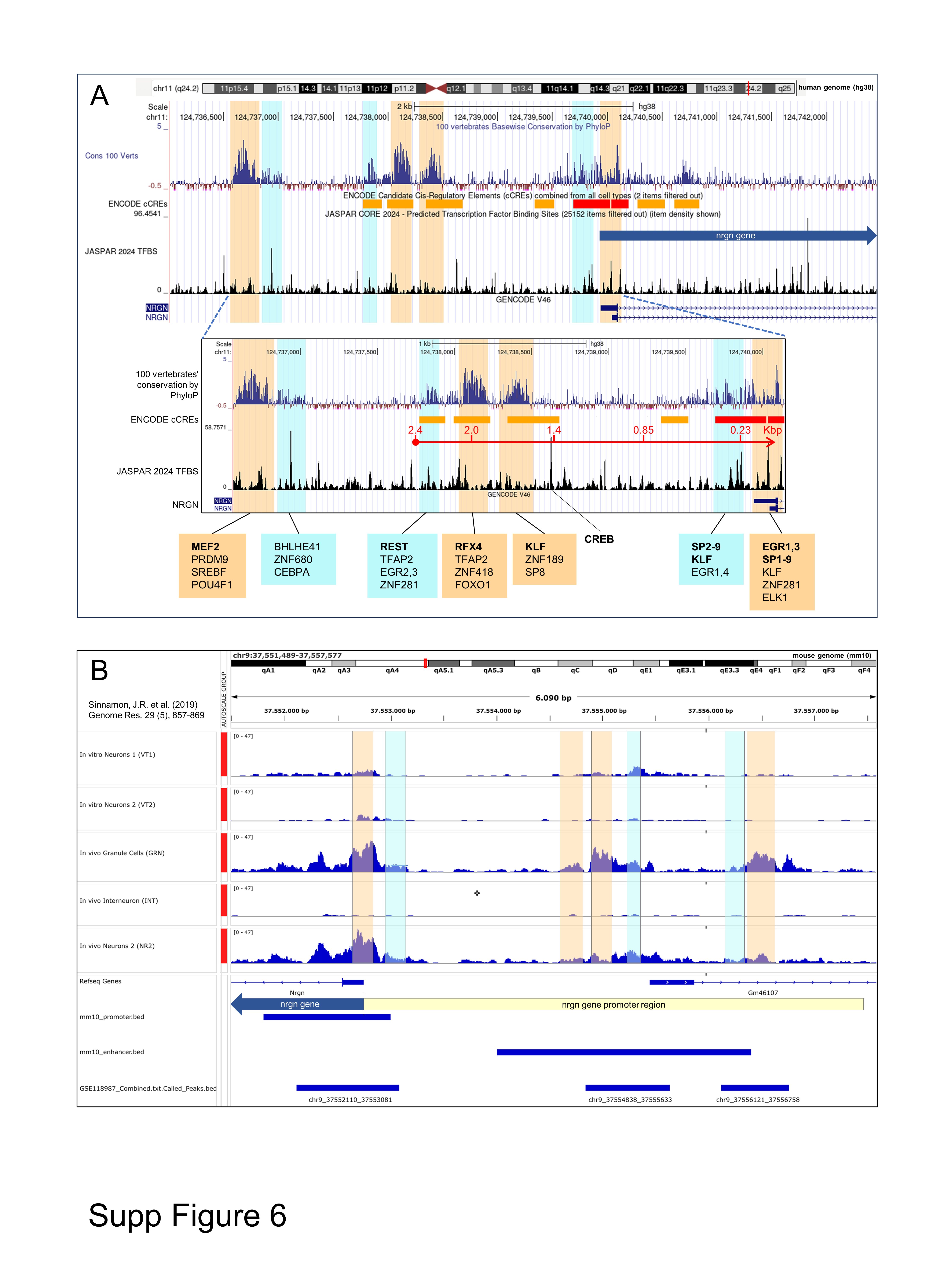

### Supp. Table 1

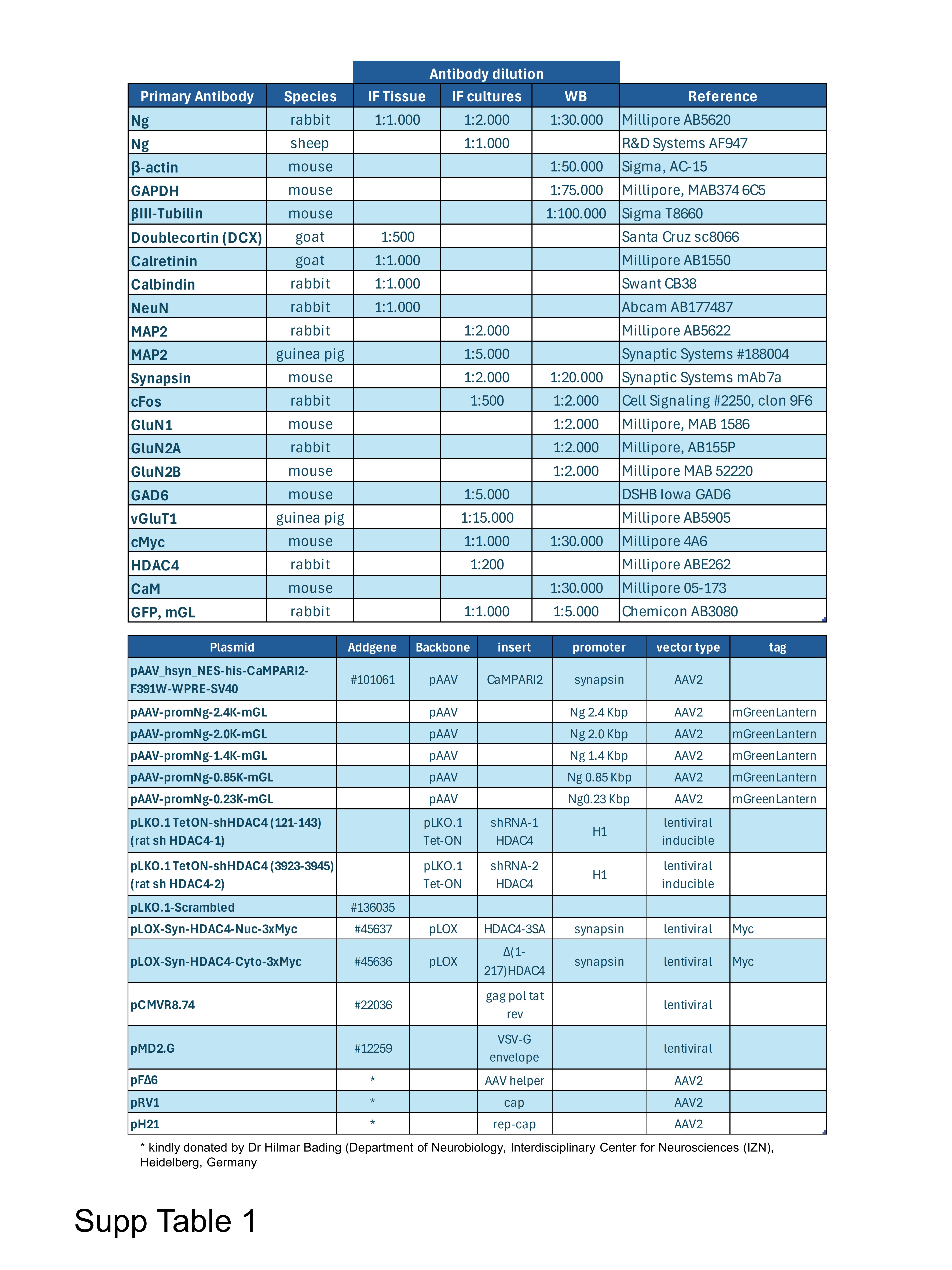
